# ConforFlux: Particle-Guided Trunk Repulsion for Diverse Protein Conformations

**DOI:** 10.64898/2026.05.16.725138

**Authors:** Shosuke Suzuki, Toshiyuki Amagasa

## Abstract

Deep learning models for protein structure prediction achieve near-experimental accuracy on individual folds, yet their default inference samples concentrate around a single dominant conformation. We introduce ConforFlux, an inference-time procedure for Boltz-2 that couples parallel structure-prediction trajectories through a pairwise C*α*-RMSD repulsion gradient on the trunk’s single and pair embeddings. The updated trunk conditions every subsequent denoising step without retraining the model. On four conformational-change categories, ConforFlux improves the per-state success rate over default Boltz-2 by 3–17 percentage points, with increased clashscores. The procedure also improves coverage on OpenFold3 and Protenix. On targets whose two reference structures were released after each model’s training cutoff, guidance improves coverage of both states on all three models. On the human dopamine transporter, ConforFlux samples inward conformations absent from 500 default samples.

## 1 Introduction

Protein function arises from transitions among conformational states rather than from a single static structure [Henzler-Wildman and Kern, 2007, Boehr et al., 2009]. Enzymes alternate between catalytically distinct conformers, membrane transporters cycle between inward- and outward-facing states, and many therapeutically relevant binding sites, including cryptic pockets, are exposed only transiently in low-population conformations. Resolving such alternate states experimentally is costly and often incomplete, so a single deposited structure typically reflects only one of several functionally relevant conformations. Computational sampling can help identify alternative structures for mechanistic studies and structure-based design.

Deep learning models for protein structure prediction now achieve near-experimental accuracy on individual folds [Jumper et al., 2021, Abramson et al., 2024, Passaro et al., 2025], yet samples from their standard inference pipelines remain concentrated around a dominant conformation for each input [Saldaño et al., 2022]. Even when a target’s alternate state is well-characterized in the literature, sampling at default settings can fail to reach it. Existing approaches broaden conformational coverage by perturbing the multiple sequence alignment (MSA), guiding the diffusion trajectory, or modifying internal representations. Other approaches train a separate model to generate conformational ensembles. We investigate whether coupling sampling trajectories through the trunk representations that condition diffusion can improve access to alternative conformations while keeping the model’s weights and inputs fixed.

We introduce ConforFlux, an inference-time procedure that couples parallel prediction trajectories through pairwise C*α*-RMSD repulsion applied to the trunk’s single and pair embeddings. In Boltz-2 [Passaro et al., 2025], these representations condition every block of the diffusion module [Abramson et al., 2024], so their updates affect subsequent denoising steps. ConforFlux adapts Particle Guidance [Corso et al., 2024] to this conditioning representation and requires neither additional training nor target-state experimental restraints.

This work makes three contributions.

- **Method**. A trunk-level particle-guidance procedure for Boltz-2 that requires no additional training (Algorithm 1). We also evaluate it on OpenFold3 and Protenix.
- **Benchmark comparison**. A controlled comparison against nine baselines spanning MSA, coordinate, embedding, and trained-ensemble interventions on the four-category ConforMix benchmark and an additional transporter set, evaluated on per-state success rate, fill ratio, and local geometry (Section 4).
- **Post-cutoff evaluation**. An evaluation of each model on targets whose two references were both released after its training cutoff, and a DAT case study over the inward, occluded and outward states of the alternating-access cycle (Sections 4.3, 4.4).

## 2 Related work

### MSA-level perturbation

Reducing or perturbing the input MSA induces AlphaFold and AlphaFold 3 to sample alternative conformations [del Alamo et al., 2022, Stein and Mchaourab, 2022, Wayment-Steele et al., 2024, Monteiro da Silva et al., 2024, Kalakoti and Wallner, 2025, 2026, Wu and Feng, 2025, Li et al., 2026a], but AlphaFold 3 and Boltz-2 have reduced the MSA’s role in favor of internal pair representations [Abramson et al., 2024, Passaro et al., 2025].

### Coordinate-level guidance

ConforMix [Richman et al., 2025] combines a classifier-guidance gradient on the diffusion variable with SMC particle resampling (Twisted Diffusion Sampler [Wu et al., 2023]), using RMSD conditioning against a baseline prediction. AF3-ReD [Ohnuki and Okazaki, 2026] adds a Gaussian repulsion between predicted atomic coordinates inside AF3 denoising. Metadiffusion [Lam et al., 2026] adds a meta-energy gradient bias to the denoised coordinates. All three perturb the denoising trajectory at coordinate level. ConforFlux back-propagates a related pairwise repulsion to the trunk representations that condition the diffusion module, then decodes through that module without modification (Section 3.2).

### Representation-level intervention

Pair representation scaling [Suzuki and Amagasa, 2026] uniformly rescales the pair representation before the Pairformer. Experiment-guided methods [Fadini et al., 2026, Li et al., 2026b, Maddipatla et al., 2026] optimize MSA cluster profiles or internal representations to fit crystallographic, cryo-EM/ET, or NMR data. ConforNets [Lee et al., 2026] trains channel-wise affine transforms of OpenFold3-preview’s pair representation before the Pairformer, jointly optimizing multiple transforms to diversify samples for each input protein. ConforFlux requires no additional training and couples the current sampling trajectories through their post-Pairformer single and pair representations.

### Particle Guidance and trained ensemble generators

Particle Guidance [Corso et al., 2024] couples multiple diffusion trajectories [Ho et al., 2020, Song et al., 2021] through a pairwise potential and is connected to Stein variational inference [Liu and Wang, 2016, Liu, 2017, D’Angelo and Fortuin, 2021]. It is instantiated on the diffusion variable in image and torsion-angle settings [Jing et al., 2022]. ConforFlux applies the coupling to the trunk representations that condition the diffusion module in AlphaFold 3 and Boltz-2 [Abramson et al., 2024, Passaro et al., 2025]. Separately trained ensemble generators include AlphaFlow [Jing et al., 2024], BioEmu [Lewis et al., 2025] and DiG [Zheng et al., 2024]. We compare with BioEmu, a model trained to approximate protein equilibrium ensembles.

## 3 Method

ConforFlux is an inference-time procedure that leaves the model’s weights and inputs unchanged (Figure 1).

**Figure 1.**
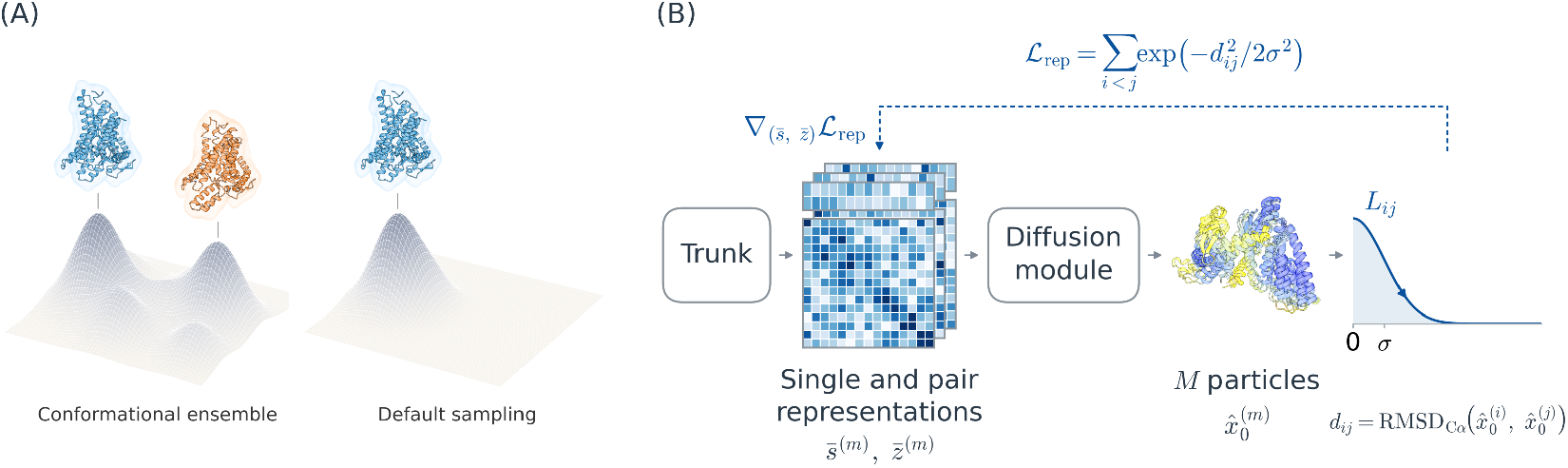
The sampling problem and the update. **(A)** Schematic comparison of a protein’s conformational ensemble and default sampling concentrated around one state. **(B)** The trunk runs once, and each trajectory receives its own copy of the single and pair representations. Pairwise C*α*-RMSD repulsion between decoded structures updates these copies by back-propagation, giving each trajectory a distinct conditioning representation.

### 3.1 The two-stage prediction pipeline

Boltz-2 [Passaro et al., 2025], OpenFold3-preview-2 [OpenFold Consortium, 2025] and Protenix [ByteDance AML AI4Science Team et al., 2025] follow the two-stage design of AlphaFold 3 [Abramson et al., 2024]. A trunk, built from an MSA module and a Pairformer stack, maps the input sequence and MSA to a single representation 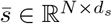 and a pair representation 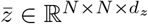, computed once per recycling iteration. A separate diffusion module *F*_*θ*_ then denoises the 3D atomic coordinates *x*_*t*_ as an all-atom point cloud, conditioned on 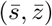 through AdaLN scale-shift [Peebles and Xie, 2023] and attention pair-bias at each of its blocks. Default sampling runs *M* reverse-diffusion trajectories independently from Gaussian noise.

### 3.2 The trunk as a guidance site

The diffusion module’s pair conditioning is built from 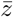 and a relative position encoding, independently of the diffusion timestep (Abramson et al., 2024, Alg. 21). This allows its computation to be amortized across the denoising trajectory [Wohlwend et al., 2024]. Updating 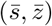 changes the conditioning of every subsequent denoising step. ConforFlux runs *M* trajectories in parallel, giving each particle its own copy (*s*^(*m*)^, *z*^(*m*)^) of the trunk output. Pairwise repulsion is evaluated on the decoded structures and back-propagated into these per-particle embeddings.

#### Algorithm 1

ConforFlux, in Boltz-2’s notation. Every line carrying (*m*) runs for all *M* particles.

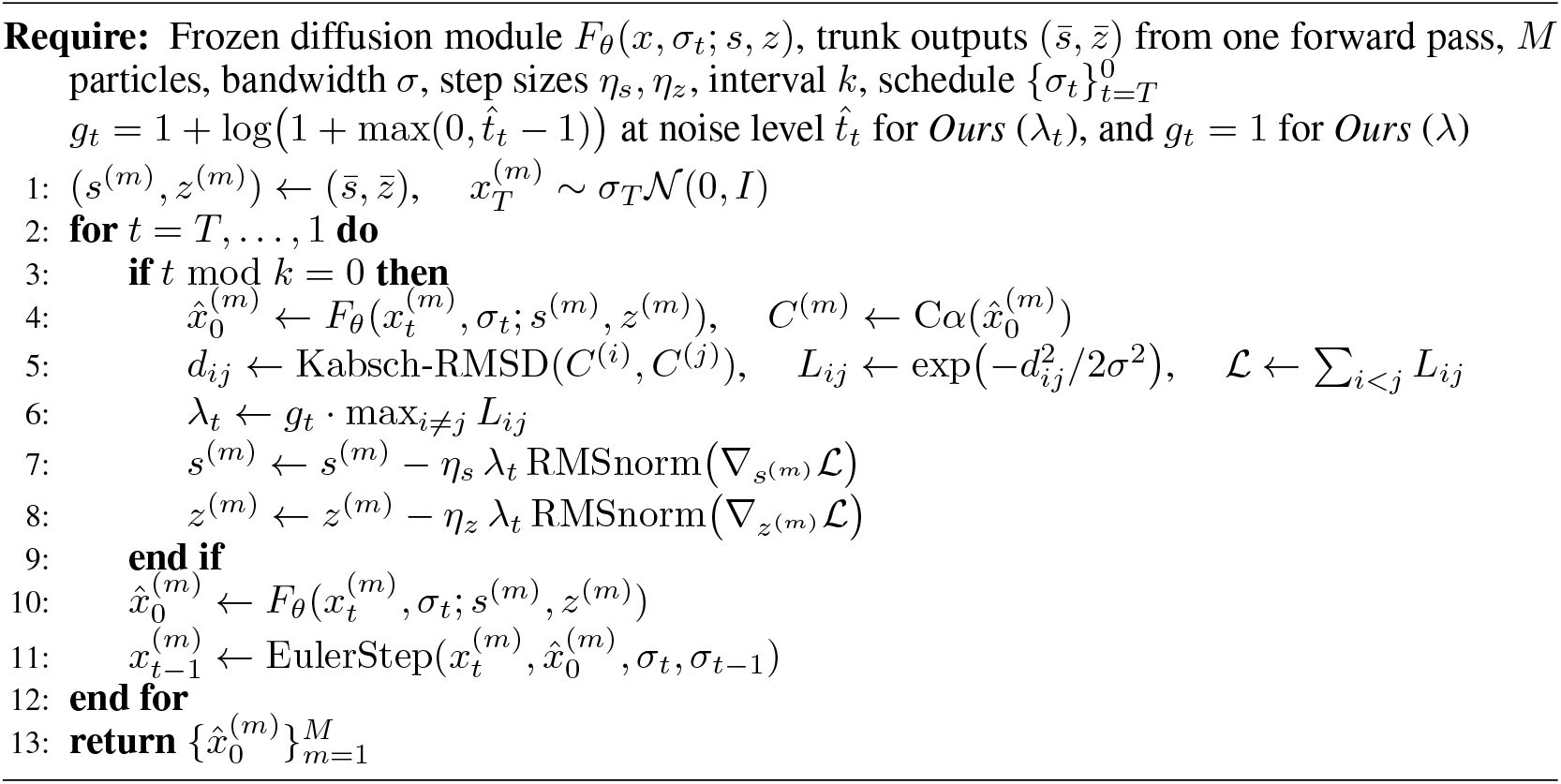

### 3.3 Pairwise C*α* repulsion across particles

At a guided step (Alg. 1, lines 4–5), we run the diffusion module independently on each particle to obtain candidate denoised structures 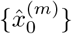, align each pair by differentiable Kabsch (SVD), and form the following RBF similarity kernel on the C*α* RMSD.

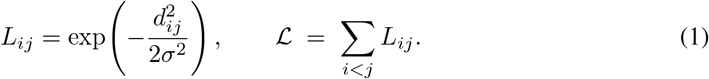

The bandwidth *σ* is held constant per particle batch, and we combine samples generated with band-widths {0.5, 1.0, 1.5, 2.0, 2.5} Å. Equation (1) is the pairwise objective of Particle Guidance [Corso et al., 2024]. We evaluate it on the diffusion module’s 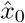 estimates, following Chung et al. [2023], Bansal et al. [2023].

### 3.4 Trunk-level descent and re-conditioning

We back-propagate ℒ to each particle’s trunk embeddings and apply an RMS-normalized descent step (Alg. 1, lines 7–8) with an adaptive scaling factor

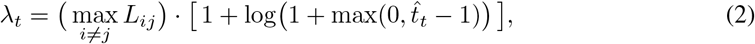

where 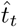 is the current noise level on the diffusion module’s EDM schedule [Karras et al., 2022]. The first factor decreases as particles separate on the scale of *σ*. The second increases with the noise level, following classifier guidance [Dhariwal and Nichol, 2021]. We use *λ*_*t*_ as the primary setting and also report *λ* = max_*i ≠ j*_ *L*_*ij*_, which omits the explicit noise-level factor while retaining adaptation to particle separation. The tables label these settings *Ours* (*λ*_*t*_) and *Ours* (*λ*).

RMS-normalization decouples step size from gradient magnitude. After each update, the AdaLN scale-shift and pair-bias projections derived from (*s*^(*m*)^, *z*^(*m*)^) are recomputed. Guidance is applied every three diffusion steps with *η*_*s*_ = *η*_*z*_ = 0.02 and *M* =5 (Appendix A).

## 4 Experiments

We evaluate on the four benchmark datasets distributed with ConforMix [Richman et al., 2025], comprising domain motion (*n*=38), membrane transporter cycling (*n*=15), cryptic pocket formation (*n*=33), and fold switching (*n*=15), each with two experimentally resolved reference conformations. We expand the transporter category with 20 pairs from Wu and Feng [2025], bringing it to 35 targets. The combined benchmark contains 121 target-category pairs. Proteins appearing in two categories are evaluated separately under each category’s protocol. Section 4.3 evaluates targets whose two references were released after each model’s training cutoff. Appendix F details the dataset construction.

All Boltz-2-based benchmark methods use the same model weights and generate 500 samples per target, spanning MSA (CF-random, AFsample3, AF-cluster), embedding (pair representation scaling), and coordinate (ConforMix) interventions. We compare with BioEmu using the same number of samples. Boltz-2 is trained on experimental structures and MD trajectory ensembles [Siebenmorgen et al., 2024, Vander Meersche et al., 2024, Mirarchi et al., 2024] with experimental-method conditioning at inference. We include both X-ray- and MD-conditioned defaults. All other Boltz-2-based methods use X-ray conditioning.

Per-state success is the fraction of reference states reached within category-specific C*α*-RMSD thresholds. RC50 is the fraction of targets for which both references are reached within half their inter-reference RMSD. Fill ratio [Kalakoti and Wallner, 2025] measures weighted bin occupancy between the references in TM-score space. We assess local geometry using MolProbity clashscore [Chen et al., 2010], focusing on interventions using Boltz-2. Appendices D and E give the thresholds and metric definitions.

### 4.1 Coverage on the four-category benchmark

ConforFlux (*λ*_*t*_) gains 3.3 to 17.1 pp over default Boltz-2 in per-state success (Table 1). Coverage improves most on transporters, with smaller gains on fold switching. The highest domain-motion success and fill ratio are obtained with *λ*, while *λ*_*t*_ gives higher transporter success and fill ratio. Both settings lead on cryptic-pocket success. BioEmu achieves the highest transporter per-state success.

**Table 1.**
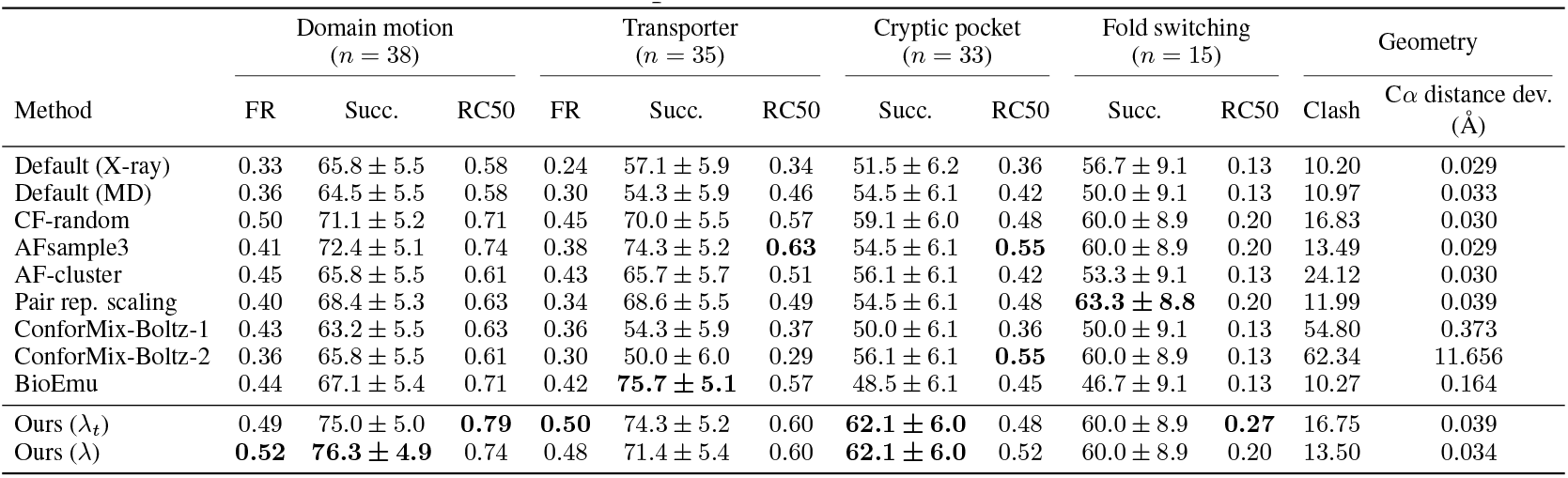
Main results by category. Per-state success rate (Succ., %), fill ratio (FR) and RC50 on the four ConforMix categories, the transporter category augmented by the 20 EGF transporter pairs of Appendix F. FR is reported for the two categories where every target is eligible for it. Clash is MolProbity’s clashscore. C*α* distance dev. is the RMS deviation of consecutive-residue C*α*–C*α* distances from 3.80 Å. The two Ours rows differ only in Eq. (2). The *λ*_*t*_ setting keeps both factors, while *λ* omits the noise-level factor. Metrics and datasets are defined in Appendices E and F. BioEmu is an external ensemble baseline. Its geometry values are reported for reference.

| Method | Domain motion<br>($n = 38$) | | | Transporter<br>($n = 35$) | | | Cryptic pocket<br>($n = 33$) | | Fold switching<br>($n = 15$) | | Geometry | |
| --- | --- | --- | --- | --- | --- | --- | --- | --- | --- | --- | --- | --- |
| | FR | Succ. | RC50 | FR | Succ. | RC50 | Succ. | RC50 | Succ. | RC50 | Clash | Cα distance dev.<br>(Å) |
| Default (X-ray) | 0.33 | 65.8 ± 5.5 | 0.58 | 0.24 | 57.1 ± 5.9 | 0.34 | 51.5 ± 6.2 | 0.36 | 56.7 ± 9.1 | 0.13 | 10.20 | 0.029 |
| Default (MD) | 0.36 | 64.5 ± 5.5 | 0.58 | 0.30 | 54.3 ± 5.9 | 0.46 | 54.5 ± 6.1 | 0.42 | 50.0 ± 9.1 | 0.13 | 10.97 | 0.033 |
| CF-random | 0.50 | 71.1 ± 5.2 | 0.71 | 0.45 | 70.0 ± 5.5 | 0.57 | 59.1 ± 6.0 | 0.48 | 60.0 ± 8.9 | 0.20 | 16.83 | 0.030 |
| AFsample3 | 0.41 | 72.4 ± 5.1 | 0.74 | 0.38 | 74.3 ± 5.2 | **0.63** | 54.5 ± 6.1 | **0.55** | 60.0 ± 8.9 | 0.20 | 13.49 | 0.029 |
| AF-cluster | 0.45 | 65.8 ± 5.5 | 0.61 | 0.43 | 65.7 ± 5.7 | 0.51 | 56.1 ± 6.1 | 0.42 | 53.3 ± 9.1 | 0.13 | 24.12 | 0.030 |
| Pair rep. scaling | 0.40 | 68.4 ± 5.3 | 0.63 | 0.34 | 68.6 ± 5.5 | 0.49 | 54.5 ± 6.1 | 0.48 | **63.3 ± 8.8** | 0.20 | 11.99 | 0.039 |
| ConforMix-Boltz-1 | 0.43 | 63.2 ± 5.5 | 0.63 | 0.36 | 54.3 ± 5.9 | 0.37 | 50.0 ± 6.1 | 0.36 | 50.0 ± 9.1 | 0.13 | 54.80 | 0.373 |
| ConforMix-Boltz-2 | 0.36 | 65.8 ± 5.5 | 0.61 | 0.30 | 50.0 ± 6.0 | 0.29 | 56.1 ± 6.1 | **0.55** | 60.0 ± 8.9 | 0.13 | 62.34 | 11.656 |
| BioEmu | 0.44 | 67.1 ± 5.4 | 0.71 | 0.42 | **75.7 ± 5.1** | 0.57 | 48.5 ± 6.1 | 0.45 | 46.7 ± 9.1 | 0.13 | 10.27 | 0.164 |
| Ours ($\lambda_t$) | 0.49 | 75.0 ± 5.0 | **0.79** | **0.50** | 74.3 ± 5.2 | 0.60 | **62.1 ± 6.0** | 0.48 | 60.0 ± 8.9 | **0.27** | 16.75 | 0.039 |
| Ours ($\lambda$) | **0.52** | **76.3 ± 4.9** | 0.74 | 0.48 | 71.4 ± 5.4 | 0.60 | **62.1 ± 6.0** | 0.52 | 60.0 ± 8.9 | 0.20 | 13.50 | 0.034 |

ConforFlux (*λ*_*t*_) reaches 28 reference states missed by default sampling, while failing to reach one state covered by default, the second reference of Q83VS8. The higher coverage comes with increased clashscores, comparable to CF-random under *λ*_*t*_ and lower under *λ* (Appendix B).

The gain over default is largest at medium difficulty, defined by default Boltz-2’s worst-case minimum RMSD (Table 2). ConforFlux also improves coverage on hard targets, while easy targets leave less room for improvement. On hard targets, *λ* gives the highest per-state success and fill ratio, and *λ*_*t*_ the highest RC50 among the methods using Boltz-2.

**Table 2.**
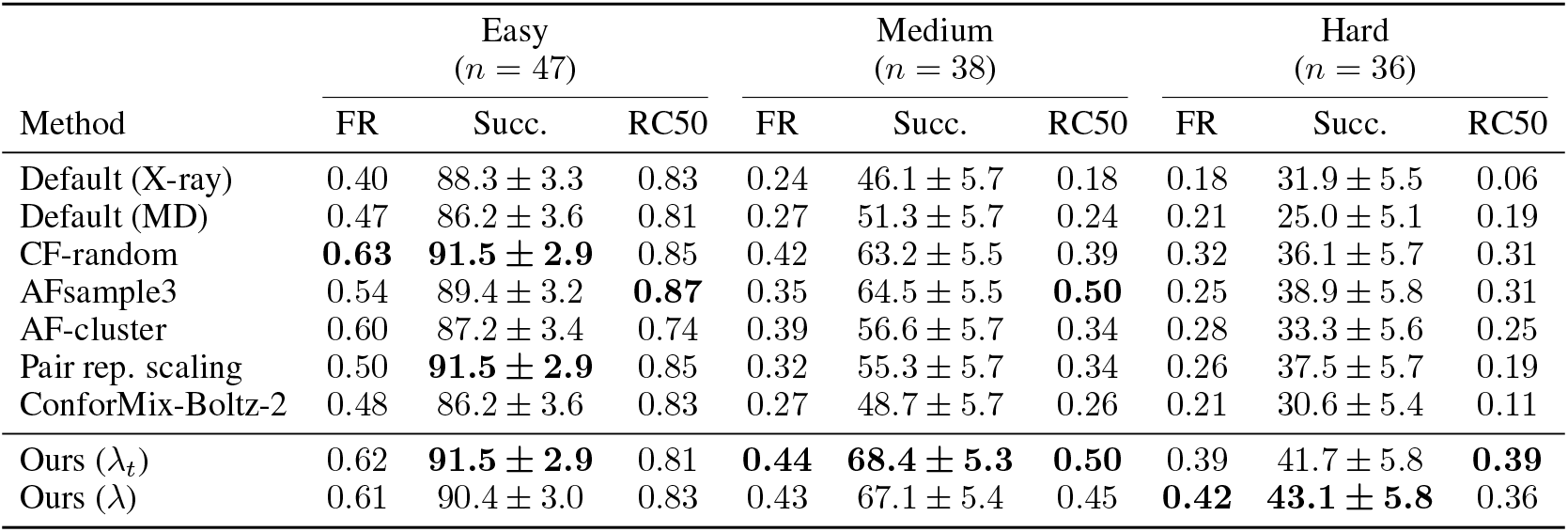
Per-state success, fill ratio and RC50 by target difficulty. Difficulty is defined by default Boltz-2’s worst-case minimum C*α* Kabsch RMSD over 500 samples, the larger of the minimum RMSDs to the two references (easy *<* 2 Å, medium 2–4 Å, hard *≥* 4 Å). FR is evaluated on the domain-motion and transporter targets, with 29, 22 and 22 in the easy, medium and hard bins, respectively. For cryptic-pocket targets, Succ. uses pocket-region C*α* Kabsch RMSD (Appendix E).

Figure 2 illustrates the effect on oligopeptidase B (O76728), bcMalT EIIC (Q63GK8) and MdfA (P0AEY8). Default sampling concentrates near one reference, while ConforFlux reaches both. Per-residue root-mean-square fluctuation (RMSF) more closely follows the inter-reference |ΔC*α*| profile on all three targets, in both amplitude and correlation.

**Figure 2.**
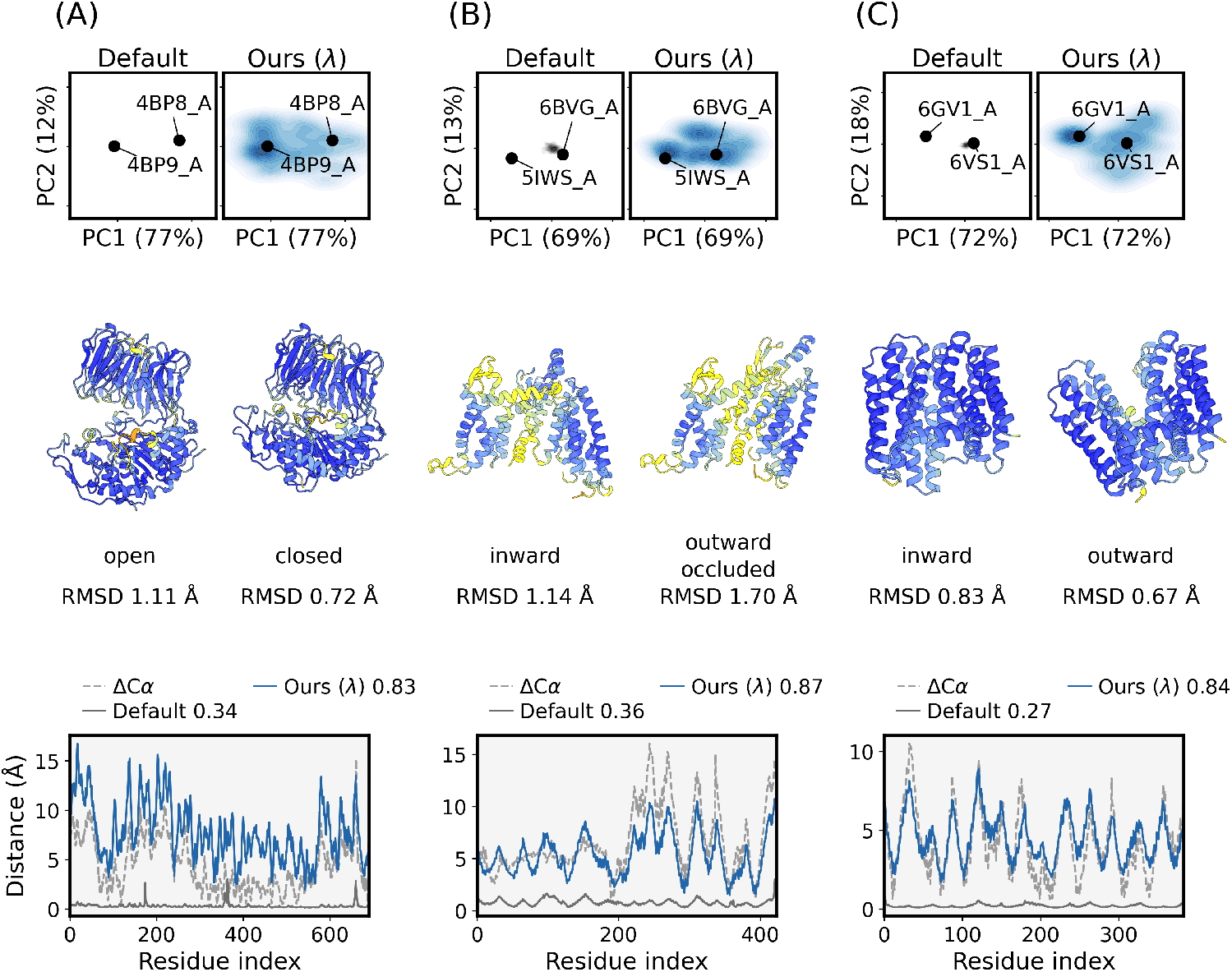
Three benchmark targets. **(A)** Oligopeptidase B (O76728, domain motion), references 4BP8_A open and 4BP9_A closed. **(B)** bcMalT EIIC (Q63GK8, membrane transporter), references 6BVG_A inward-facing and 5IWS_A outward-facing occluded. **(C)** MdfA (P0AEY8, membrane transporter), references 6VS1_A inward-facing and 6GV1_A outward open. **Top**, PCA of pairwise C*α* distances. **Middle**, the closest ConforFlux prediction to each reference among the 500 samples. **Bottom**, the per-residue RMSF against the inter-reference |ΔC*α*| profile, with the Pearson correlation shown in each panel. The panels show ConforFlux with *λ* at the settings in Table 7. The analysis uses the same residue set as the RMSD evaluation. PCA uses samples retained by the pLDDT filter of Appendix D. RMSF and numerical evaluations use all 500 samples.

### 4.2 Transfer to OpenFold3 and Protenix

We apply the same guidance to OpenFold3-preview-2 and Protenix, whose trunk representations condition a separate diffusion module. Both settings of Eq. (2) are evaluated on the main benchmark, using 100 samples per target for both guided and default sampling (Table 3). Under *λ*_*t*_, per-state success increases on domain motion, transporters and fold switching on both models, and RC50 increases in every category. Cryptic-pocket success changes little under this setting, while MolProbity scores change by at most 0.26. Omitting the noise-level factor produces less ensemble spread and a smaller increase in clashscore, with category-dependent changes in success.

**Table 3.** ConforFlux on OpenFold3 and Protenix. Categories and metrics as in Table 1, with 100 samples per target for both guided and default sampling. Both models use the guidance settings in Table 7.

| Model | Method | Domain motion<br>($n = 38$) | | | Transporter<br>($n = 35$) | | | Cryptic pocket<br>($n = 33$) | | Fold switching<br>($n = 15$) | |
| --- | --- | --- | --- | --- | --- | --- | --- | --- | --- | --- | --- |
|  |  | FR | Succ. | RC50 | FR | Succ. | RC50 | Succ. | RC50 | Succ. | RC50 |
| OpenFold3-p2 | Default | 0.18 | 60.5 ± 5.6 | 0.45 | 0.14 | 67.1 ± 5.6 | 0.49 | 45.5 ± 6.1 | 0.21 | 53.3 ± 9.1 | 0.07 |
| | Ours ($\lambda_t$) | 0.31 | 71.1 ± 5.2 | **0.74** | **0.34** | 74.3 ± 5.2 | **0.63** | 45.5 ± 6.1 | **0.36** | **56.7 ± 9.1** | **0.13** |
| | Ours ($\lambda$) | **0.32** | **72.4 ± 5.1** | **0.74** | 0.33 | **75.7 ± 5.1** | **0.63** | **54.5 ± 6.1** | **0.36** | 50.0 ± 9.1 | **0.13** |
| Protenix | Default | 0.21 | 61.8 ± 5.6 | 0.42 | 0.19 | 67.1 ± 5.6 | 0.51 | 51.5 ± 6.1 | 0.27 | 40.0 ± 8.9 | 0.13 |
| | Ours ($\lambda_t$) | 0.30 | 71.1 ± 5.2 | **0.74** | **0.35** | **74.3 ± 5.2** | **0.71** | 53.0 ± 6.1 | 0.42 | **50.0 ± 9.1** | **0.20** |
| | Ours ($\lambda$) | **0.31** | **72.4 ± 5.1** | 0.71 | 0.34 | **74.3 ± 5.2** | **0.71** | **56.1 ± 6.1** | **0.48** | 43.3 ± 9.0 | **0.20** |

Both models have an earlier training cutoff than Boltz-2. Section 4.3 evaluates each on targets whose two reference structures were released after its training cutoff.

### 4.3 Coverage beyond each model’s training cutoff

We evaluate each model on targets whose two reference structures were both released after its training cutoff, which is 2023-06-01 for Boltz-2 and 2021-09-30 for OpenFold3-preview-2 and Protenix. The selected sets contain 12 targets for Boltz-2 and 20 for the other two models (Appendix F). Within each model’s set, we compare default sampling and ConforFlux using RC50 and alternate-state RMSD (Table 4).

**Table 4.**
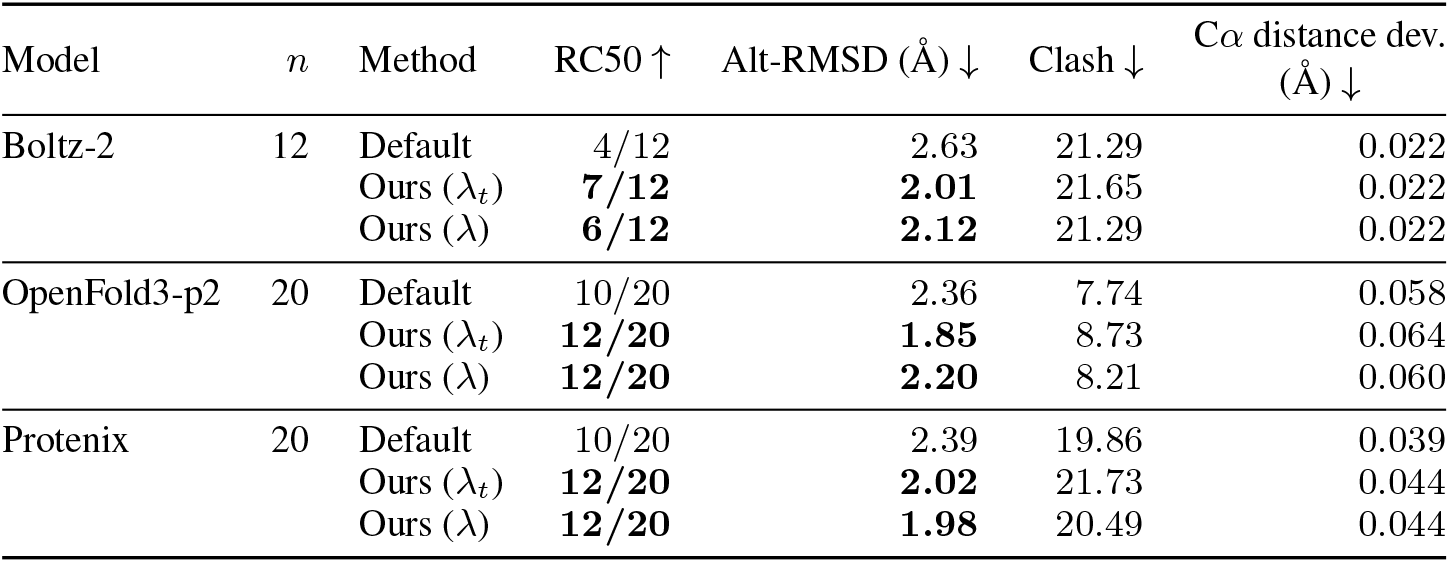
Coverage beyond each model’s training cutoff. Targets whose two reference structures were both released after that model’s cutoff, selected as described in Appendix F. RC50 is a count of targets here rather than a fraction, and Alt-RMSD the worst-case minimum C*α* RMSD averaged over them. Boltz-2 draws 500 samples per target and the other two 100. The three blocks have different targets and different cutoffs.

ConforFlux (*λ*_*t*_) improves RC50 on all three models and reduces alternate-state RMSD by 0.37 to 0.63 Å. Figure 3 illustrates the gain on OCT1. Under *λ*, one fewer Boltz-2 target meets RC50 than under *λ*_*t*_, while the counts are equal on the other models. Clashscores are lower under *λ*. This evaluation tests coverage of newly released structures. Homologs in related conformations may occur in the training data.

**Figure 3.**
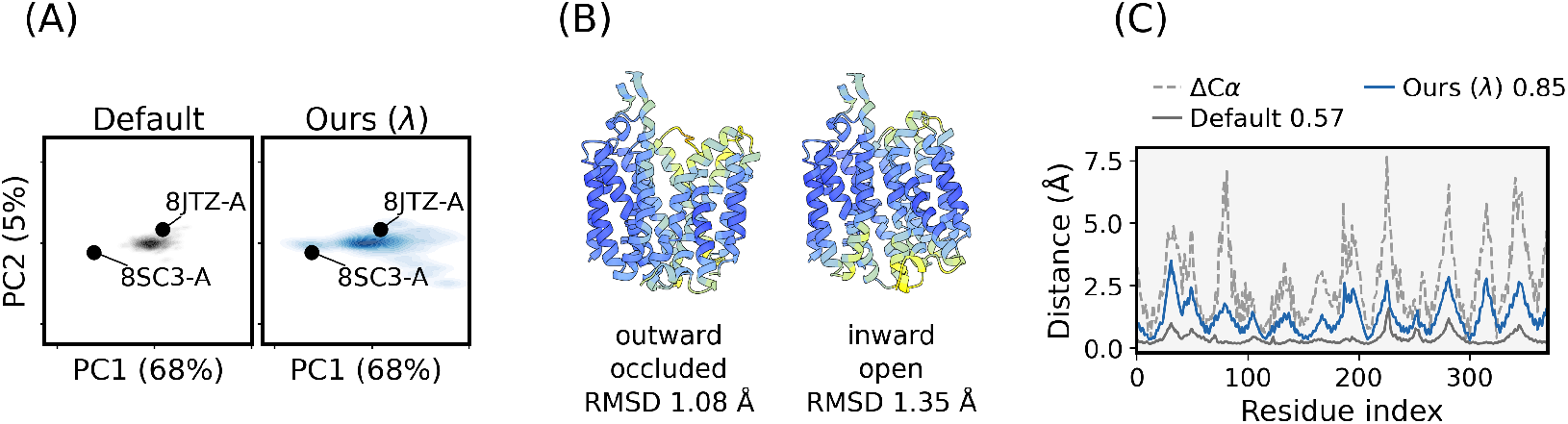
A post-cutoff transporter. OCT1 (O15245), references 8JTZ outward-facing partially occluded and 8SC3 inward-open. **(A)** PCA of pairwise C*α* distances. **(B)** The closest prediction to each reference. **(C)** Per-residue RMSF compared with the dashed inter-reference |ΔC*α*| profile, with the Pearson correlation shown in each panel. PCA uses samples retained by the pLDDT filter of Appendix D. RMSF and numerical evaluations use all 500 samples.

### 4.4 Mechanistic case study on the dopamine transporter

We run a case study on the human dopamine transporter (DAT, UniProt Q01959). Li et al. [2024] solved five cryo-EM structures of DAT (PDB 8Y2C, 8Y2D, 8Y2E, 8Y2F, 8Y2G) that together capture the outward-facing, occluded, and inward-facing states of the alternating-access cycle, all post-cutoff. DAT is not in the post-cutoff transporter set. In the pairwise-distance PCA of Figure 4, default predictions cluster in an intermediate region between the occluded and outward references, while ConforFlux samples extend to all three states. Assigning each prediction to the best-matching state at a 0.3 Å RMSD margin, default reaches the inward state in 0*/*500 samples and ConforFlux in 43*/*500.

**Figure 4.**
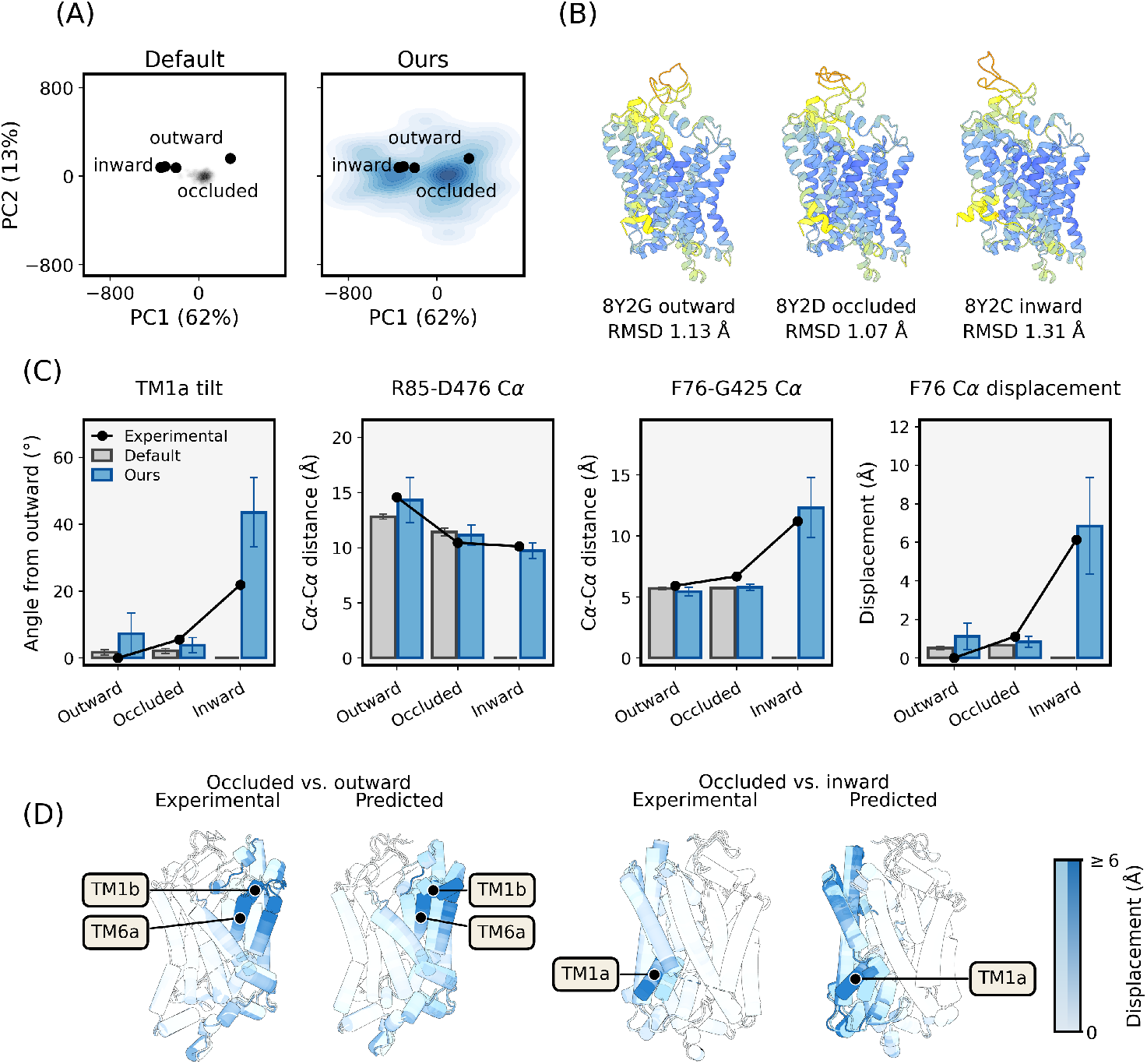
DAT case study. Each ensemble contains 500 samples generated at the settings of Table 7. **(A)** Pairwise-distance PCA with the five experimental references shown as black dots. **(B)** The closest ConforFlux prediction to each state, with its reference RMSD below. **(C)** Means and standard deviations of four structural measurements by assigned state. Class sizes are reported in Table 11. **(D)** Experimental and predicted changes from occluded to outward and inward states. Residue color and opacity indicate displacement on a shared scale across all four panels. PCA uses samples retained by the pLDDT filter of Appendix D. State assignments and structural measurements use all 500 samples.

We assess the predictions assigned to the inward-facing state using four C*α*-only measurements of the outward-to-inward transition [Li et al., 2024]. R85–D476 separation, F76–G425 cavity opening, and F76 displacement are close to the experimental inward values, while TM1a tilt is larger (Figure 4C and Appendix K). Default predictions retain a closed cytoplasmic cavity. These measurements support the inward assignment, with an overestimate of the helix tilt.

### 4.5 Intervention site

We compare trunk guidance with two controls on the main benchmark at 100 samples per target (Table 5). The noise control uses isotropic trunk updates matched to the guidance step norm. The coordinate control applies the same repulsion to denoised coordinates, with *η*_*x*_ swept over 0.02, 0.2, and 2.0.

**Table 5.** Intervention site. 121 target–category pairs, 100 samples each. Matched noise replaces the repulsion gradient with an isotropic step of the same norm at the same site. The coordinate rows keep the repulsion and move it to the denoised coordinates at step size *η*_*x*_. The four geometry columns are per-sample means. The last column reports the share of samples with at least one consecutive-residue C*α*–C*α* distance differing from 3.80 Å by more than 1.0 Å.

| Method | Coverage |  |  | Geometry |  |  |  |
| --- | --- | --- | --- | --- | --- | --- | --- |
| | FR | Succ. (%) | RC50 | MolProbity | Clash | Rama out. (%) | Abnormal $C\alpha$ distances (%) |
| Default | 0.190 | 55.0 | 0.322 | 1.51 | 10.2 | 0.04 | 0.4 |
| Ours (trunk) | **0.307** | **61.6** | **0.496** | 1.69 | 16.5 | 0.07 | 4.9 |
| trunk, matched noise | 0.137 | 52.1 | 0.223 | 1.50 | 10.1 | 0.04 | 0.3 |
| coordinates, $\eta_x = 0.02$ | 0.142 | 51.2 | 0.248 | 1.52 | 10.3 | 0.08 | 6.5 |
| coordinates, $\eta_x = 0.2$ | 0.245 | 57.4 | 0.339 | 2.26 | 21.0 | 2.48 | 57.4 |
| coordinates, $\eta_x = 2.0$ | 0.274 | 61.2 | 0.455 | 1.88 | 16.0 | 0.76 | 24.5 |

Matched noise reduces coverage relative to default sampling, supporting the importance of the repulsion direction. Coordinate repulsion at *η*_*x*_ = 0.02 also reduces coverage. A larger coordinate step recovers similar per-state success to trunk guidance but lower fill ratio and RC50, with more Ramachandran outliers. Here the C*α*-only objective moves only C*α* atoms under coordinate guidance, whereas the trunk update is decoded into a whole structure by the denoiser (Figure 5).

**Figure 5.**
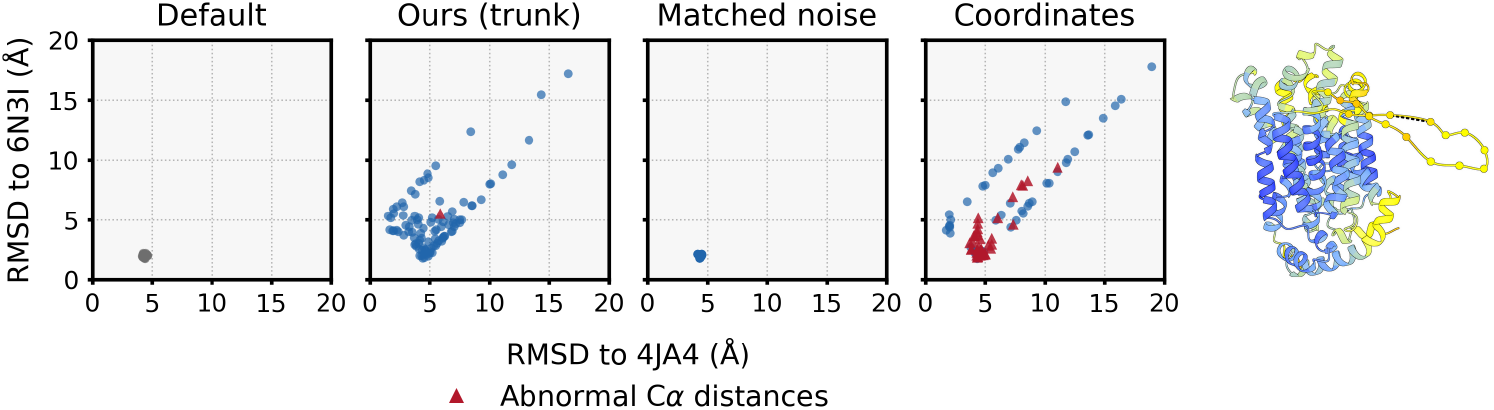
Intervention site on one target. P0AGF4, 100 samples per method, each at its C*α* RMSD to the two references (4JA4 chain C, 6N3I chain A). The coordinate panel is *η*_*x*_ = 2.0. The right panel shows the coordinate-guided sample with the most abnormal consecutive-residue C*α*–C*α* distances, as defined in Table 5. Spheres mark the affected C*α* atoms, and the dashed line marks the longest distance.

## 5 Discussion

A trunk update influences subsequent denoising steps through their conditioning, while the diffusion module generates coordinates. The transfer results support this guidance site across the three tested models, and the matched-noise control supports the importance of the repulsion direction at a matched update norm on Boltz-2. Trunk and coordinate step sizes are not directly comparable because the updates act in different spaces. At similar per-state success in the tested sweep, coordinate guidance has lower fill ratio and RC50 and more Ramachandran outliers than trunk guidance.

On Boltz-2, gains peak at medium difficulty and persist on hard targets. Recovering reference states missed by matched default ensembles shows access to additional states at a fixed budget. RMSF follows reference displacements more closely on the highlighted targets, and DAT measurements support inward gate opening. Thus, the broader sampling includes experimentally observed changes without specifying a target conformation.

Fold-switching gains remain modest on Boltz-2. Lower aggregate confidence is consistent with a shift toward samples farther from the references, since confidence at a given distance from the nearer reference remains similar across settings (Appendix C). Confidence alone cannot distinguish unobserved conformations from prediction errors far from both references, and state frequencies under repulsion should not be interpreted as equilibrium populations.

## 6 Conclusion and limitations

ConforFlux couples parallel sampling trajectories through the trunk’s conditioning representations. Repulsion between the current predictions provides a structured update direction without specifying a target conformation. On Boltz-2, norm-matched noise does not reproduce the coverage gain. Across three models, ConforFlux recovers experimental reference states missed by default sampling at the same sample budget, without additional training.

On the Boltz-2 benchmark, fold switching remains difficult, with modest gains over default sampling and success comparable to the other inference-time baselines. Broader coverage also comes with higher clashscores and a larger fraction of samples with abnormal consecutive-residue C*α* distances (Appendix B and Table 5). ConforFlux uses more diffusion time and peak memory than ConforMix. Gradient checkpointing trades additional diffusion time for lower peak memory (Appendix G).

## Computational resources

This research used computational resources of Pegasus at the Center for Computational Sciences, University of Tsukuba, through the Multidisciplinary Cooperative Research Program, and of SQUID [Date et al., 2023] at the D3 Center, The University of Osaka, through the Project for Nurturing Student Competing with the World.

## Code availability

Source code is available at github.com/suzuki-2001/conforflux.

## Ethics statement

This work uses only publicly released structures and benchmarks. It involves no human or animal subjects and uses no sensitive personal data.

## APPENDIX

### A Experimental details and hyperparameter settings

#### A.1 Base model configuration

The main benchmark uses Boltz-2 [Passaro et al., 2025], with ConforMix-Boltz-1 on Boltz-1 [Wohlwend et al., 2024] and BioEmu as a separately trained ensemble model. Transfer experiments use OpenFold3-preview-2 and Protenix. Both ConforMix baselines set the step scale to 1.0 [Richman et al., 2025]. Other parameters follow the respective base-model defaults (Table 6).

**Table 6.**
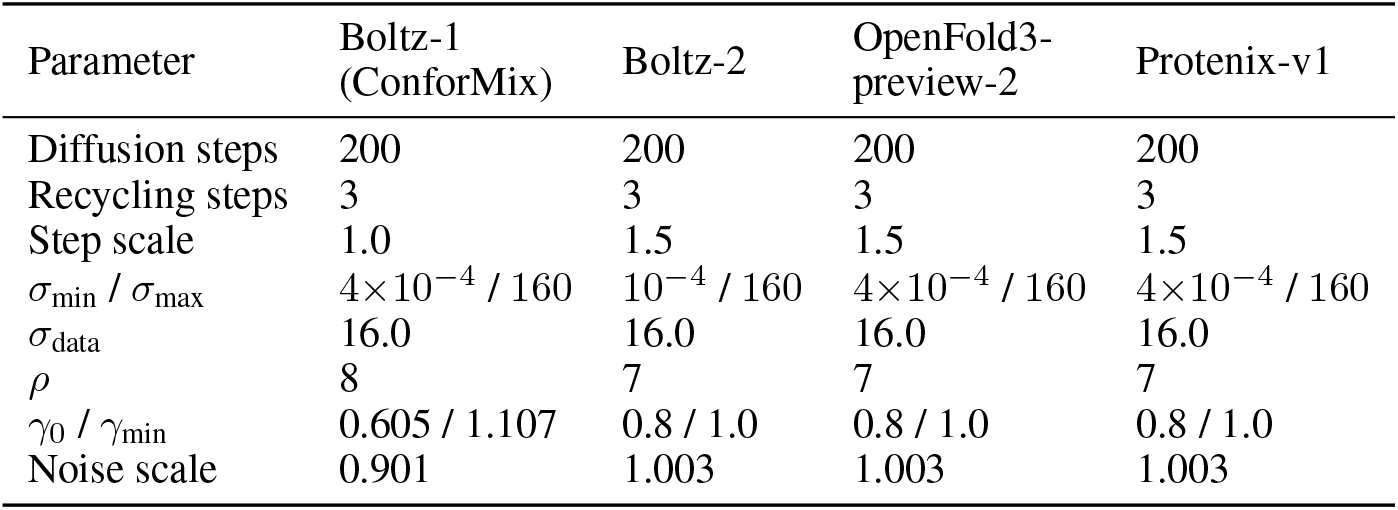
Base model inference parameters.

| Parameter | Boltz-1<br>(ConforMix) | Boltz-2 | OpenFold3-preview-2 | Protenix-v1 |
| --- | --- | --- | --- | --- |
| Diffusion steps | 200 | 200 | 200 | 200 |
| Recycling steps | 3 | 3 | 3 | 3 |
| Step scale | 1.0 | 1.5 | 1.5 | 1.5 |
| $\sigma_{\min} / \sigma_{\max}$ | $4 \times 10^{-4} / 160$ | $10^{-4} / 160$ | $4 \times 10^{-4} / 160$ | $4 \times 10^{-4} / 160$ |
| $\sigma_{\text{data}}$ | 16.0 | 16.0 | 16.0 | 16.0 |
| $\rho$ | 8 | 7 | 7 | 7 |
| $\gamma_0 / \gamma_{\min}$ | 0.605 / 1.107 | 0.8 / 1.0 | 0.8 / 1.0 | 0.8 / 1.0 |
| Noise scale | 0.901 | 1.003 | 1.003 | 1.003 |

**Table 7.**
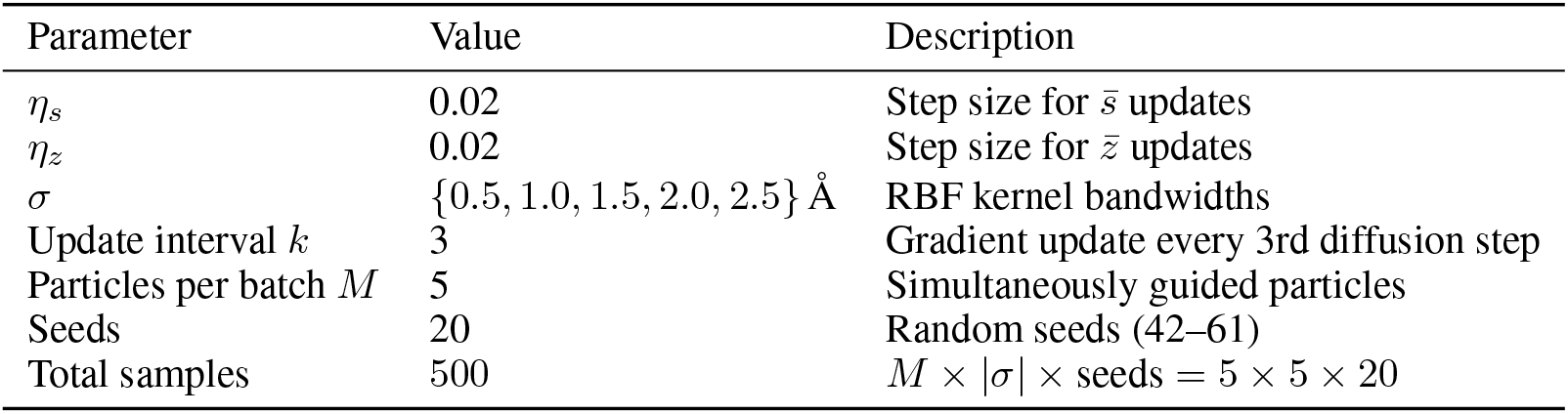
ConforFlux guidance hyperparameters for the 500-sample Boltz-2 benchmark.

#### A.2 ConforFlux guidance

Embedding updates are RMS-normalized and scaled by the maximum pairwise kernel value in Eq. (2). The primary setting additionally uses the noise-level factor 1 + log(1 + max(0, 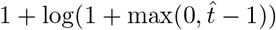)), where 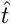is the current noise level on the diffusion module’s EDM schedule [Karras et al., 2022]. The kernel bandwidth *σ* is held constant per particle batch and swept across the values in Table 7. The Boltz-2 benchmark uses 20 seeds and 500 samples per target. Transfer and intervention-site experiments use the same guidance settings with 100 samples per target.

#### A.3 Baseline methods

Table 8 summarizes the configurations used for the nine baselines. All methods generate 500 samples per target on the same MSA, computed with the ColabFold MMseqs2 server [Mirdita et al., 2022].

**Table 8.** Baseline method configurations. The base model is Boltz-2 unless otherwise noted.

| Method | Key parameters | Sampling configuration |
| --- | --- | --- |
| Default (X-ray) | Seeds 42–51 | 10 seeds × 50 samples = 500. Boltz-2 inference with X-ray experimental-method conditioning. |
| Default (MD) | Seeds 42–51 | Same MSA and seeds as Default (X-ray). Boltz-2 inference with MD experimental-method conditioning. |
| CF-random | Depths<br>$\{2, 4, 8, 16, 32, 64, 128\}$ | 100 runs × 5 samples = 500. Each run subsamples $d-1$ homologs, evenly across the seven depths. |
| AFsample3 | Mask rate 0.40 | 100 mask seeds × 5 diffusion samples = 500. Boltz-2 port of the AF3 setting reported by Kalakoti and Wallner [2026]. |
| AF-cluster | DBSCAN with ε-sweep | 500 samples spread evenly over up to five clusters, at the $\varepsilon$ maximizing the cluster count, following Wayment-Steele et al. [2024]. |
| Pair representation scaling | $\beta \in \{\pm 0.15, \pm 0.30, \pm 0.45, \pm 0.60, \pm 0.75\}$ | 10 values × 50 samples = 500. The pair representation is scaled by $(1+\beta)$ before the Pairformer [Suzuki and Amagasa, 2026]. |
| ConforMix-Boltz-1 | $\alpha = 15$, RMSD range 0–20 Å, secondary-structure mask | 25 RMSD targets × 20 samples = 500, the official implementation of Richman et al. [2025] on our MSA. |
| ConforMix-Boltz-2 | Same as Boltz-1 | The same sampler over the Boltz-2 denoiser and data pipeline (Appendix I). |
| BioEmu | – | 500 samples per target from the released model [Lewis et al., 2025], on the same MSA as the other baselines. |

#### A.4 Software

Boltz-2 is used at v2.2.1, OpenFold3-preview-2 at checkpoint `openfold3-p2-155k`, and Protenix at `protenix_base_default_v1.0.0`, the checkpoint whose training data ends 2021-09-30. The ConforMix-Boltz-1 baseline uses the bundled Boltz-1 v0.3.0 distributed with the ConforMix reference implementation (https://github.com/drorlab/conformix). BioEmu uses the released bioemu-v1.1 model with base seed 42 and sample filtering disabled. Structure visualizations are rendered with UCSF ChimeraX [Meng et al., 2023].

### B Structural quality

We assess local geometry using MolProbity [Chen et al., 2010] and C*α* distance deviation (Table 9), focusing on interventions using Boltz-2. BioEmu learns approximate equilibrium ensembles and generates backbone structures through a different model [Lewis et al., 2025]. These local metrics do not assess ensemble populations.

**Table 9.** Structural quality. MolProbity scores and C*α* distance deviations over the 121 target-category pairs, with 500 samples generated per pair. Distance deviation is the RMS deviation of consecutive-residue C*α*–C*α* distances from 3.80 Å. ConforMix is reported before filtering, after physicality filtering, and after both physicality and pLDDT filtering.

| Method | Clashscore<br>(↓) | $C\alpha$ distance<br>deviation (Å, ↓) | Ramachandran (%) | |
| --- | --- | --- | --- | --- |
|  |  |  | fav (↑) | out (↓) |
| Default (X-ray) | **10.20** | **0.029** | 98.81 | 0.04 |
| Default (MD) | 10.97 | 0.033 | **98.99** | **0.03** |
| CF-random | 16.83 | 0.030 | 98.80 | 0.05 |
| AFsample3 | 13.49 | **0.029** | 98.85 | 0.04 |
| AF-cluster | 24.12 | 0.030 | 98.76 | 0.06 |
| Pair rep. scaling | 11.99 | 0.039 | 98.79 | 0.05 |
| ConforMix-Boltz-1 (unfiltered) | 54.80 | 0.373 | 96.47 | 0.73 |
| Physicality | 41.44 | 0.072 | 97.09 | 0.44 |
| Physicality + pLDDT | 37.07 | 0.070 | 97.47 | 0.31 |
| ConforMix-Boltz-2 (unfiltered) | 62.34 | 11.656 | 94.10 | 1.37 |
| Physicality | 29.24 | 0.048 | 96.81 | 0.46 |
| Physicality + pLDDT | 29.04 | 0.048 | 96.83 | 0.45 |
| BioEmu | 10.27 | 0.164 | 93.18 | 1.34 |
| Ours ($\lambda_t$) | 16.75 | 0.039 | 98.68 | 0.07 |
| Ours ($\lambda$) | 13.50 | 0.034 | 98.75 | 0.05 |

ConforMix’s released pipeline applies a physicality filter based on sequential C*α*–C*α* and C–N distances and the closest heavy-atom pair [Richman et al., 2025]. We report unfiltered results, physicality filtering alone, and physicality combined with the ten-residue pLDDT-window filter at a decrease threshold of 0.2. Following the released filter implementation, the pLDDT reference is the per-residue minimum over the 20 samples generated with target RMSD zero. These samples bypass pLDDT rejection. Filtering reduces bond deviations and clashes, with clashscores remaining above default sampling.

Clashscore rises from default’s 10.20 to 16.75 under ConforFlux (*λ*_*t*_), comparable to CF-random. Omitting the noise-level factor lowers clashscore to 13.50. Ramachandran outliers and C*α* distance deviations also increase relative to default sampling, with smaller changes under *λ*.

### C Boltz-2 confidence under guidance

The confidence analysis uses the 121 main benchmark target–category pairs and 12 additional EGF pairs. All targets are monomers, so Boltz-2’s per-sample confidence reduces to 0.8 · pLDDT + 0.2 · pTM [Passaro et al., 2025]. Figure 6 compares each target’s median confidence with default’s. Across these 133 pairs, median confidence decreases for 91% under *λ* and 92% under *λ*_*t*_, with median decreases of 0.008 and 0.014, respectively. Decreases tend to be larger on targets with higher default confidence.

**Figure 6.**
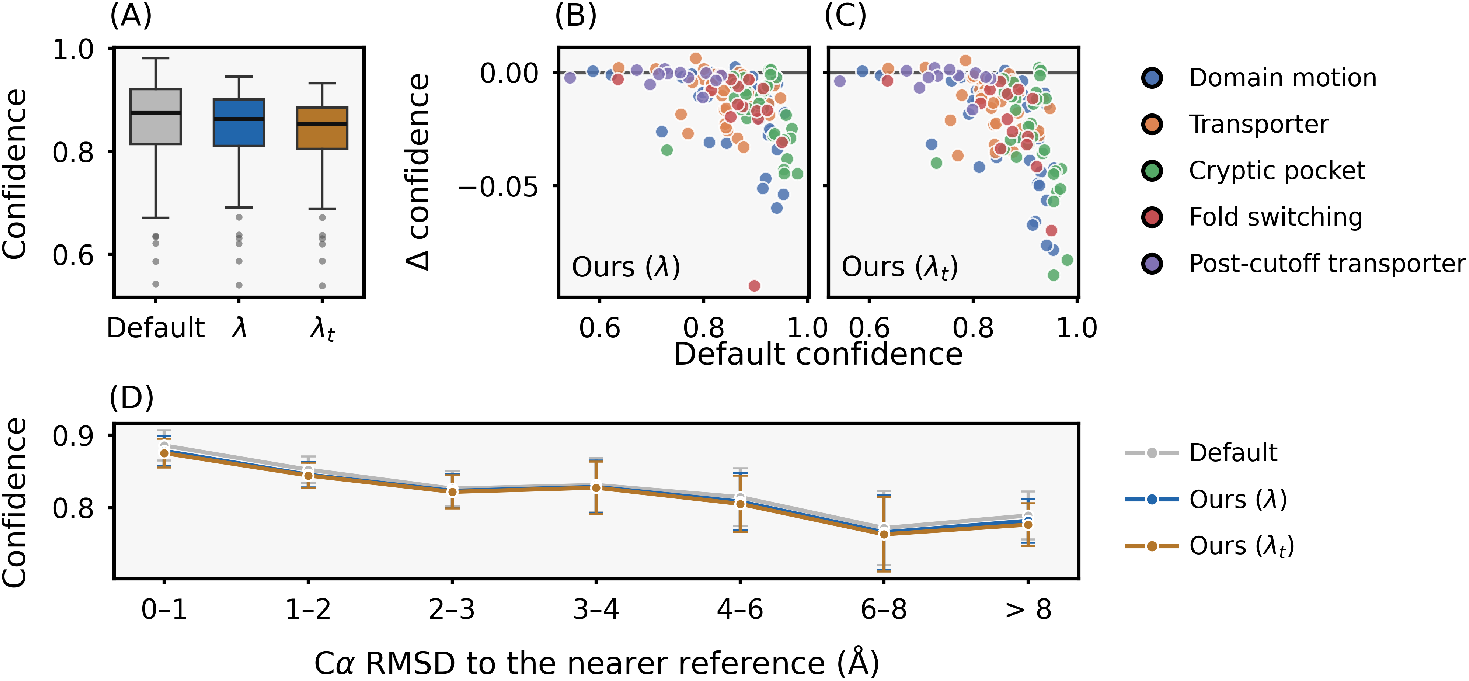
Boltz-2 confidence under guidance. **(A)** Distribution of median confidence across targets. **(B, C)** Each target’s median confidence compared with default’s for the two settings of Eq. (2). **(D)** Per-sample confidence against the C*α* RMSD to the nearer reference, averaged within each target and then across targets with samples from all three methods in the bin, with 95% intervals.

Per sample, confidence falls with distance from a reference, from 0.886 within 1 Å to 0.772 at 6 to 8 Å under default sampling. The two guided settings differ from that curve by at most 0.013 in every bin (Figure 6D). The confidence–distance relation is therefore similar across settings, while guidance shifts the distribution of distances. Default places 33.9% of its samples within 1 Å of a reference and 4.5% beyond 8 Å, against 10.1% and 19.1% under *λ*_*t*_.

### D Evaluation details

#### RMSD computation

We use the two reference structures of Richman et al. [2025] per target and compute C*α* Kabsch RMSD [Kabsch, 1976] on the sequence-aligned residue set. For cryptic pocket, RMSD is measured on the BioEmu pocket residues after superposition on the BioEmu alignment residues [Lewis et al., 2025], which coincide with the pocket residues on all 33 targets.

#### Per-residue RMSF

For the RMSF profiles in Figures 2 and 3, we superpose each sample and the second reference onto the first reference by C*α* Kabsch alignment. At each matched residue, RMSF is the root-mean-square deviation from the ensemble mean coordinate. The reference profile is the displacement between the superposed references.

#### Per-state success rate

Each (target, reference) pair contributes a binary success indicator **1** {min_*i*_ RMSD(*s*_*i*_, *r*) ≤ *τ*} over the generated ensemble. The category mean averages these indicators across all pairs. We generate 500 samples per target for the Boltz-2 benchmark and 100 for transfer and intervention-site experiments. Q16539 is excluded from cryptic pocket because the BioEmu dataset provides no pocket annotation for it, giving *N* =33 targets.

#### Category-specific thresholds

Membrane transporter uses *τ* = 2.0 Å on the full structure, following the EGF benchmark [Wu and Feng, 2025]. Domain motion also uses *τ* = 2.0 Å on the full structure, reduced from BioEmu’s 3 Å to use the same threshold as transporters. The median inter-reference C*α* RMSD is 5.20 Å for domain motion and 4.46 Å for transporters. Fold switching uses *τ* = 3.0 Å on the full structure following Wayment-Steele et al. [2024].

Cryptic pocket uses *τ* = 1.0 Å on the BioEmu pocket region [Lewis et al., 2025], compared with BioEmu’s 1.5 Å threshold. BioEmu averages per-atom distances over four backbone atoms, while we take the root mean square over C*α*.

#### Bootstrap standard deviation

Success rates are reported as the mean ± bootstrap standard deviation over (target, state) pairs. We resample these pairs with replacement over 10^5^ seeded replicates and take the standard deviation of the replicate means.

#### PCA visualization

PCA in Figures 2, 3 and 4 is fitted to samples retained by a windowed-pLDDT filter adapted from ConforMix [Richman et al., 2025]. The per-residue reference is the minimum pLDDT over the 500 default samples, which are all retained. A guided sample is excluded if its mean pLDDT decrease from this reference exceeds 0.05 in any ten-residue window. The ConforMix quality analysis uses a threshold of 0.2 (Appendix B). RMSF and numerical evaluations use all 500 generated samples in these case studies.

### E Evaluation metrics

We report four metrics.

#### Worst-case minimum RMSD

For a target with two reference structures *r*_1_, *r*_2_ and a generated set {*s*_*i*_}, we report

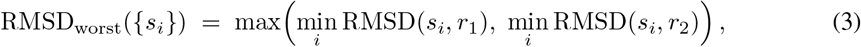

Lower values indicate closer coverage of both experimentally resolved reference states.

#### Fill ratio (FR)

Introduced by Kalakoti and Wallner [2025], FR measures occupancy along the segment between two references in TM-score space. TM-align [Zhang and Skolnick, 2005, 2004] scores each sample against both references using the fixed *d*_0_ = 3.5 Å specified in that paper. The point (TM(*s*_*i*_, *r*_1_), TM(*s*_*i*_, *r*_2_)) is projected orthogonally onto the segment joining the two references. We divide this segment into *N* = 100 equal bins weighted by *w*(*k*) = 1 + 16 (*k/*(*N* − 1) − 1*/*2)^2^, which gives greater weight to the ends of the segment. The fill ratio is

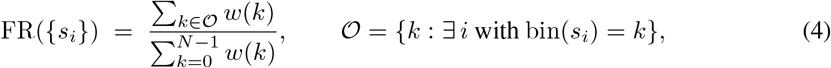

FR = 1 is an ensemble occupying every bin, FR = 0 one with no sample between the references. FR counts occupied bins, independently of the number of samples in each bin.

FR is restricted to targets with *T*_ref_ *<* 0.95, retaining all 38 domain-motion targets and all 35 transporters. Above this threshold, the references are too similar for the projection to distinguish coverage reliably. FR is reported for domain motion and transporters. Cryptic-pocket changes are local, whereas FR scores the whole structure, and fold-switching references have different folds. The projection measures occupancy along the segment and discards distance from it.

#### RC50

A target meets RC50 when its worst-case minimum RMSD is at most half the inter-reference RMSD [Richman et al., 2025]. We report the fraction of targets meeting this criterion, which requires coverage of both references relative to the size of their conformational change. For cryptic pockets, both RMSDs are measured on the annotated pocket region.

#### Clashscore

Clashscore is MolProbity’s rate of pairwise atomic overlaps per sample [Chen et al., 2010], from the clashscore2 routine of Williams et al. [2018] as implemented in Phenix CCTBX.

### F Datasets

We evaluate on the four benchmark datasets distributed with ConforMix [Richman et al., 2025] and the EGF transporter set of Wu and Feng [2025].

#### ConforMix benchmark

The four categories are domain motion (*n* = 38, BioEmu test split [Lewis et al., 2025] and OC23 [Kalakoti and Wallner, 2025]), membrane transporter cycling (*n* = 15, IOMemP/TP16 [Xie and Huang, 2024], after ConforMix excluded one of the 16 original pairs), cryptic pocket formation (*n* = 33, BioEmu cryptic-pocket set after excluding Q16539 with no pocket annotation), and fold switching (*n* = 15, following Porter and Looger [2018]).

#### EGF transporter set

We use 32 of the original 37 pairs, excluding slc39, mdfA, AAC3, PF0708, and murJ for overlap with ConforMix transporters. Table 1 adds 20 EGF pairs to the transporter category. The confidence analysis includes all 32 EGF pairs.

For Section 4.3, we select eligible pairs from all 32 EGF pairs and the ConforMix benchmarks, requiring both reference structures to have been released after the evaluated model’s cutoff. At the Boltz-2 cutoff of 2023-06-01 [Passaro et al., 2025], 19 EGF pairs have both references within cutoff, 12 have both after cutoff, and GlyT1 spans the boundary. The eligible pairs include SLC1A5, whose references were released in 2024. At the 2021-09-30 cutoff of OpenFold3-preview-2 and Protenix, 20 pairs qualify, comprising 17 EGF transporters and three ConforMix domain-motion targets (A0QTT2, P03047, Q83VS8). Both evaluations exclude P23975, whose metadata lists the same structure twice, and P0A0J7, which is not an EGF reference pair. Q01959 (DAT) is evaluated separately in the case study of Section 4.4.

### G Computational overhead

Figure 7 profiles diffusion time and peak GPU memory for default, ConforFlux, and ConforFlux with gradient checkpointing on the ConforMix benchmark. ConforFlux guidance raises mean diffusion time from 25 to 56 s, a factor of 2.2, and mean peak GPU memory from 3.9 to 21.8 GB, a factor of 5.6. Without gradient checkpointing, two targets exceeding 800 residues run out of memory. Gradient checkpointing removes those failures, with mean peak memory of 10.8 GB and mean diffusion time of 70 s, both 2.8 times default.

**Figure 7.**
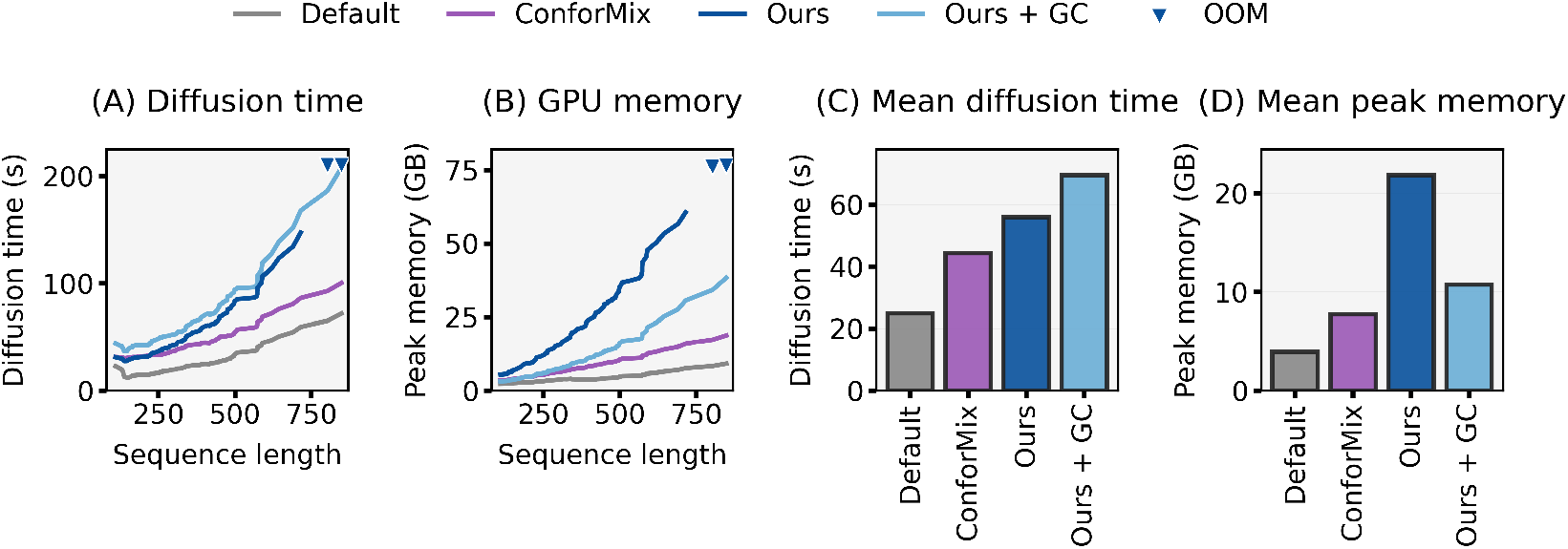
Computational overhead of ConforFlux guidance. All measurements use *M* =5 particles on a single NVIDIA H100 (80 GB). **(A)** Diffusion time versus sequence length. **(B)** Peak GPU memory versus sequence length. Downward triangles mark out-of-memory failures. **(C)** Benchmark-wide mean diffusion time. **(D)** Benchmark-wide mean peak GPU memory. Reported wall times cover the diffusion loop and, for ConforMix, generation of the reference prediction.

### H Hyperparameter sensitivity and embedding ablation

We vary one factor at a time around the main benchmark settings of Section 3. These use step size *η*_*s*_ = *η*_*z*_ = 0.02 on both trunk embeddings, *M* = 5 particles, guidance every *k* = 3 diffusion steps, the bandwidth swept over *σ* ∈ {0.5, 1.0, 1.5, 2.0, 2.5} and both factors of Eq. (2) enabled. The sweep runs on all 38 domain-motion targets. We adjust the number of seeds to generate 100 samples per target in every condition. The baseline uses the first 100 samples from the benchmark run. Metrics follow Appendices D and E.

Step size, bandwidth, and adaptive scaling have the largest effects (Figure 8). Large step sizes or bandwidths worsen both coverage and geometry. At *σ* = 8 Å, fill ratio falls from 0.290 to 0.151 and MolProbity rises from 1.62 to 2.06. The main settings combine samples generated with several kernel bandwidths in one ensemble. Omitting scaling by the maximum pairwise kernel value worsens all three metrics. Removing the noise-level factor improves them, including a 0.013 increase in fill ratio and a 0.08 Å reduction in worst-case RMSD.

**Figure 8.**
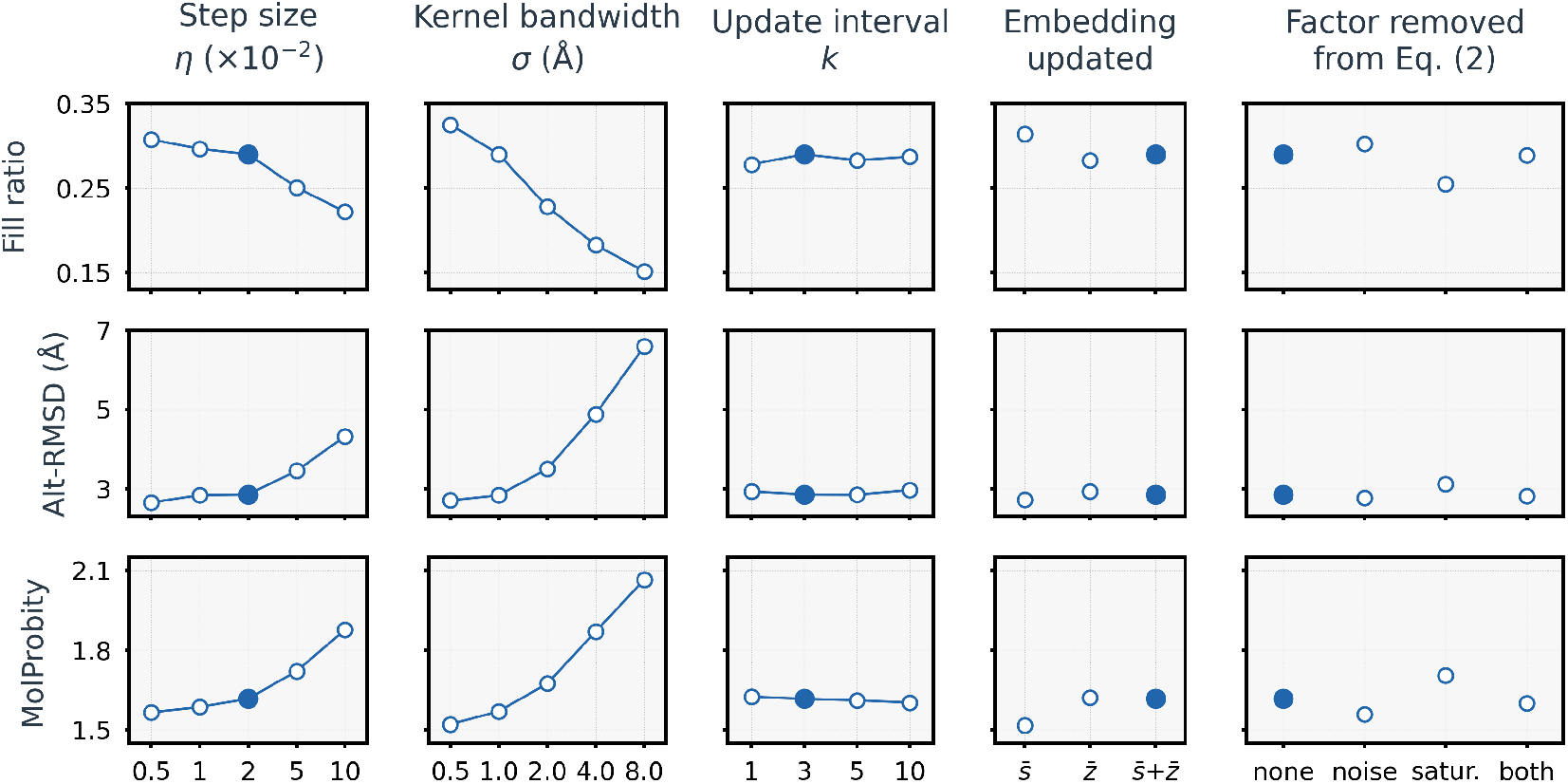
ConforFlux ablation. Fill ratio, the worst-case minimum C*α* RMSD and MolProbity over each factor in turn, on all 38 domain-motion targets at 100 samples per target. The filled marker is the configuration of Table 7. Lines join ordered levels only.

Update interval and the choice of single, pair, or combined embedding updates produce smaller changes across the tested settings. The main experiments use combined 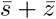 updates. Pair-only perturbations also broaden sampling, consistent with Suzuki and Amagasa [2026]. Disabling both scaling factors gives values close to those obtained with the main settings. Particle count is held at *M* = 5 throughout.

### I ConforMix reproduction

We port ConforMix [Richman et al., 2025] from Boltz-1 to Boltz-2 (configuration in Table 8). On P0205, the example target from the official repository, the Boltz-2 port and the Boltz-1 reference both produce RMSD distributions broader than their respective defaults (Figure 9). Unfiltered and physically filtered geometry results are reported in Table 9.

**Figure 9.**
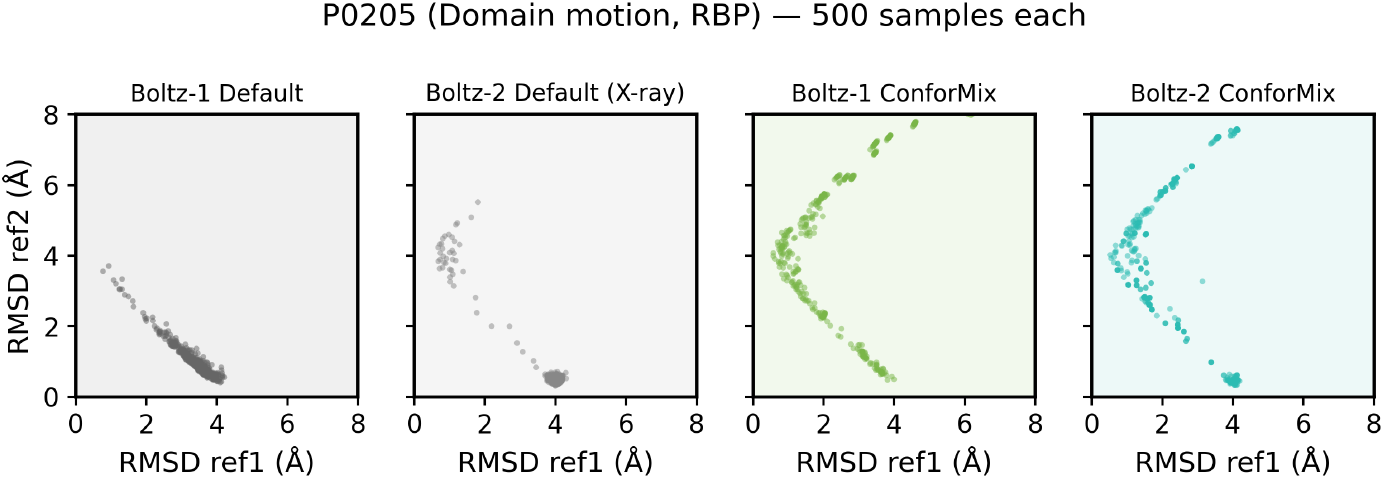
ConforMix reproduction on P0205 (RBP), the example target from the official repository. Sequence-aligned C*α* Kabsch RMSD scatter against the two reference conformations.

**Figure 10.**
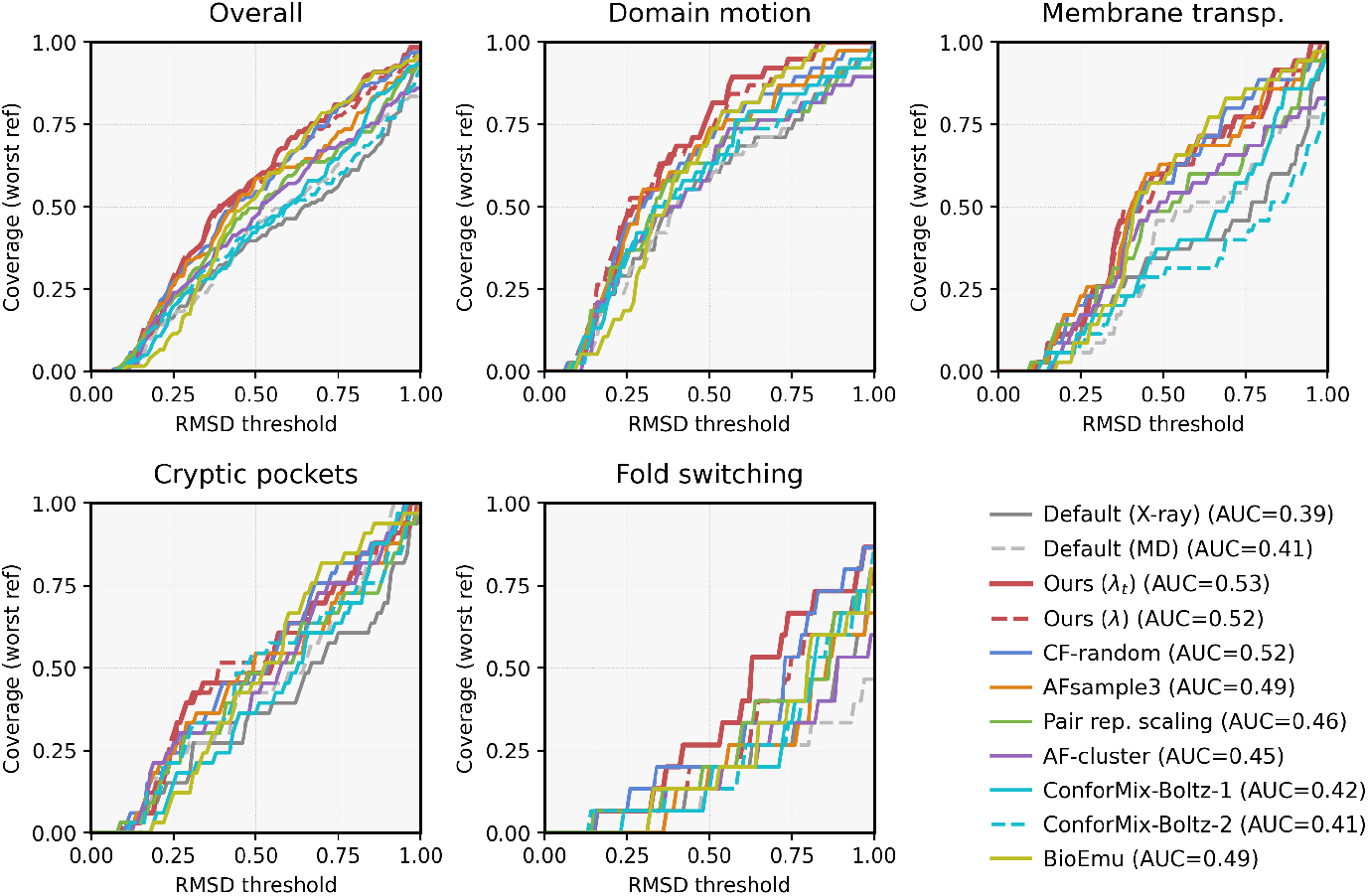
RMSD coverage curves (worst-case reference) for all methods, shown overall and per category. The shared legend reports overall AUC values.

### J RMSD coverage curves

Coverage at threshold *t* is the fraction of targets for which at least one sample reaches the worst-case reference within *t* × RMSD_ref_, where RMSD_ref_ is the distance between the two references. The transporter panel matches the Transporter category in Table 1, and Q16539 is excluded from the cryptic-pocket panel (Appendix D). ConforFlux has the highest AUC overall and in three of the four categories, second to CF-random on cryptic pockets.

### K Measurement details for the DAT case study

#### K.1 Structures, samples, and superposition

Li et al. [2024] resolved five cryo-EM structures of the human dopamine transporter (hDAT, UniProt Q01959). 8Y2G is outward-facing at 2.8 Å with methylphenidate and 8Y2D occluded at 2.8 Å with dopamine. The other three are inward-facing and lie within 0.58 Å global C*α* RMSD of one another and are treated as one inward-facing state. These are 8Y2C apo at 3.2 Å, 8Y2E with benztropine and 8Y2F with GBR12909 at 3.0 Å. All five were deposited 2024-01-25 and released 2024-08-14, after the Boltz-2 training cutoff of 2023-06-01 [Passaro et al., 2025].

We generate 500 samples each with default Boltz-2 and ConforFlux, at the configuration of Table 7. Each prediction is superimposed onto 8Y2G on the scaffold C*α* atoms (93 residues in TM3, TM4, TM8 and TM9 from the 8Y2G secondary-structure annotation) by Kabsch superposition [Kabsch, 1976] as implemented in Biopython [Cock et al., 2009].

#### K.2 Structural measurements

Table 10 defines four C*α*-only structural measurements, each tracking a rearrangement described by Li et al. [2024]. For each helix, tilt is the angle between the first principal axes fitted to its C*α* coordinates in the prediction and in 8Y2G, oriented N → C. Displacements are Euclidean distances between the prediction and 8Y2G. Both are measured after scaffold superposition. Intramolecular distances require no superposition.

**Table 10.** The four structural measurements and the rearrangement each measures.

| Measurement | Residues | Rearrangement |
| --- | --- | --- |
| TM1a tilt (°) | 66–78 | TM1a tilts by 25° from occluded to inward, opening the cytoplasmic cavity |
| R85–D476 distance (Å) | R85, D476 | their Cα separation falls from 15 to 10 Å into the occluded state, where the two form a salt bridge |
| F76–G425 distance (Å) | F76, G425 | the cytoplasmic cavity, between the residue on TM1a that moves most and one lining the inward-state inhibitor site |
| F76 displacement (Å) | F76 | the same TM1a motion, at one residue |

**Table 11.** Mean structural measurements by method and assigned state. State labels are assignments to the best-matching state at a 0.3 Å margin in global C*α* Kabsch RMSD. *n* is the class size. All nine non-empty method–class combinations are reported.

| Method | Class | $n$ | R85–D476 (Å) | F76–G425 (Å) | F76 disp. (Å) | TM1a tilt (°) |
| --- | --- | --- | --- | --- | --- | --- |
| Default | Outward | 244 | 12.81 | 5.68 | 0.51 | 1.7 |
| Default | Occluded/Outward | 249 | 12.36 | 5.70 | 0.57 | 1.6 |
| Default | Occluded | 7 | 11.41 | 5.71 | 0.67 | 2.1 |
| ConforFlux | Outward | 236 | 14.32 | 5.43 | 1.11 | 7.3 |
| ConforFlux | Occluded/Outward | 61 | 12.63 | 5.69 | 0.78 | 3.6 |
| ConforFlux | Occluded | 19 | 11.13 | 5.78 | 0.84 | 3.9 |
| ConforFlux | Inward/Occluded | 139 | 10.15 | 7.17 | 1.93 | 17.0 |
| ConforFlux | Inward/Outward | 2 | 11.00 | 6.57 | 2.58 | 14.8 |
| ConforFlux | Inward | 43 | 9.71 | 12.30 | 6.84 | 43.5 |
| Cryo-EM | 8Y2G outward | – | 14.58 | 5.91 | 0.00 | 0.0 |
| Cryo-EM | 8Y2D occluded | – | 10.44 | 6.68 | 1.11 | 5.5 |
| Cryo-EM | 8Y2C/E/F inward (mean) | – | 10.10 | 11.21 | 6.12 | 21.9 |

TM1b and TM6a tilts, which seal the extracellular pathway on the transition to occluded, and D79 C*α* displacement are defined the same way and reported in Figure 11.

**Figure 11.**
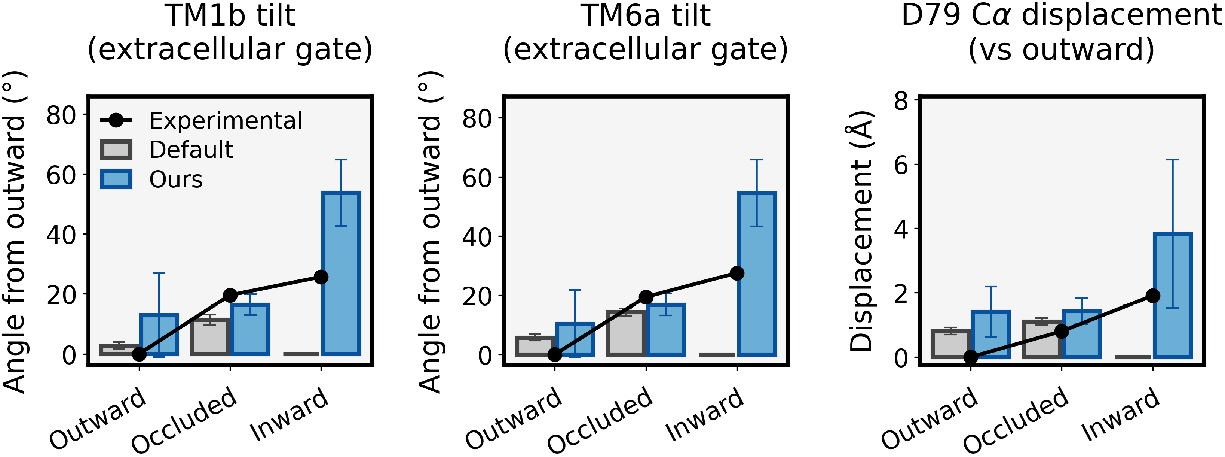
Supplementary DAT structural measurements. TM1b and TM6a tilts, which seal the extracellular pathway, move with TM1a, and D79 C*α* displacement agrees with experiment within one standard deviation.

**Figure 12.**
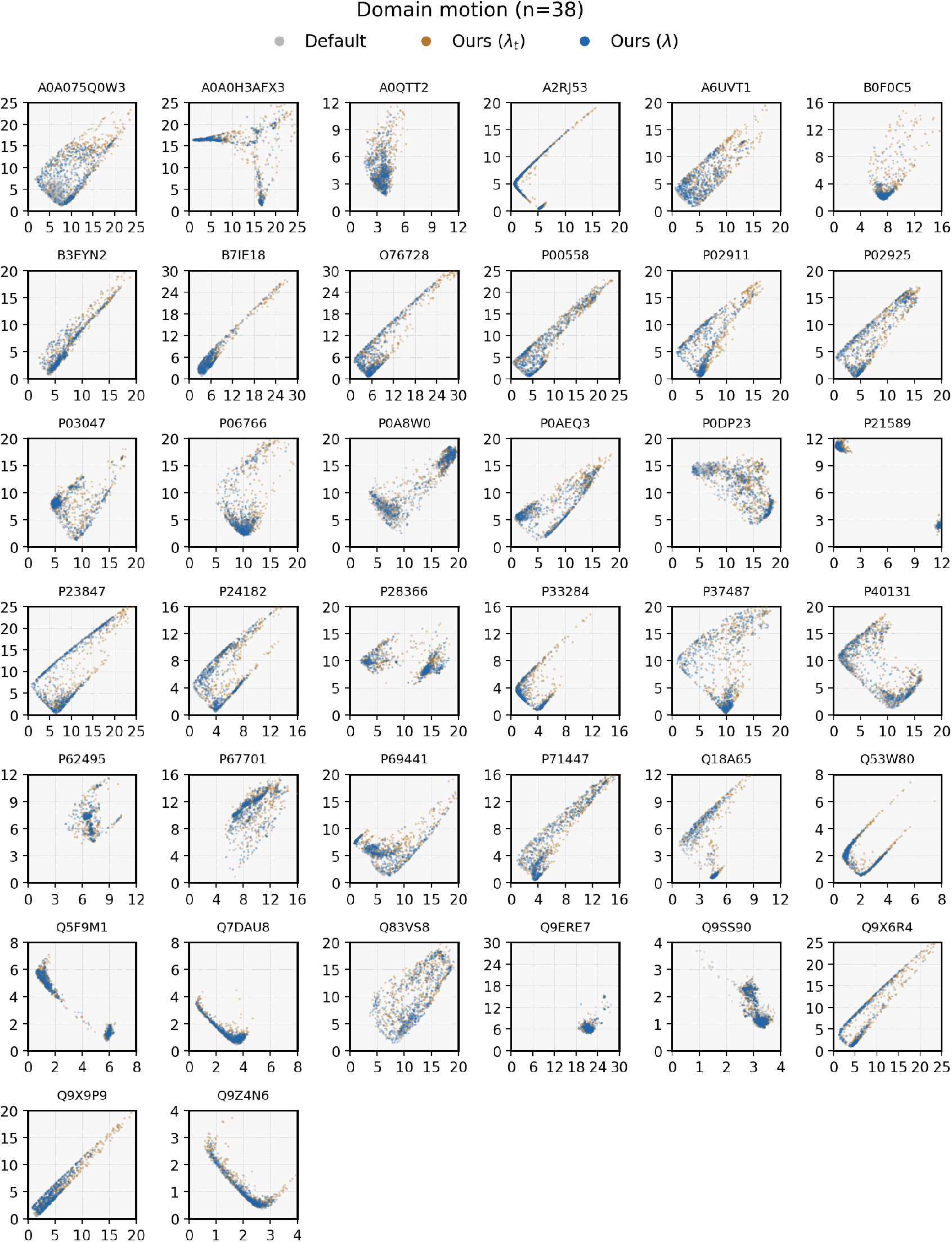
**Per-target 2D RMSD scatter for domain motion** (*n*=38).

**Figure 13.**
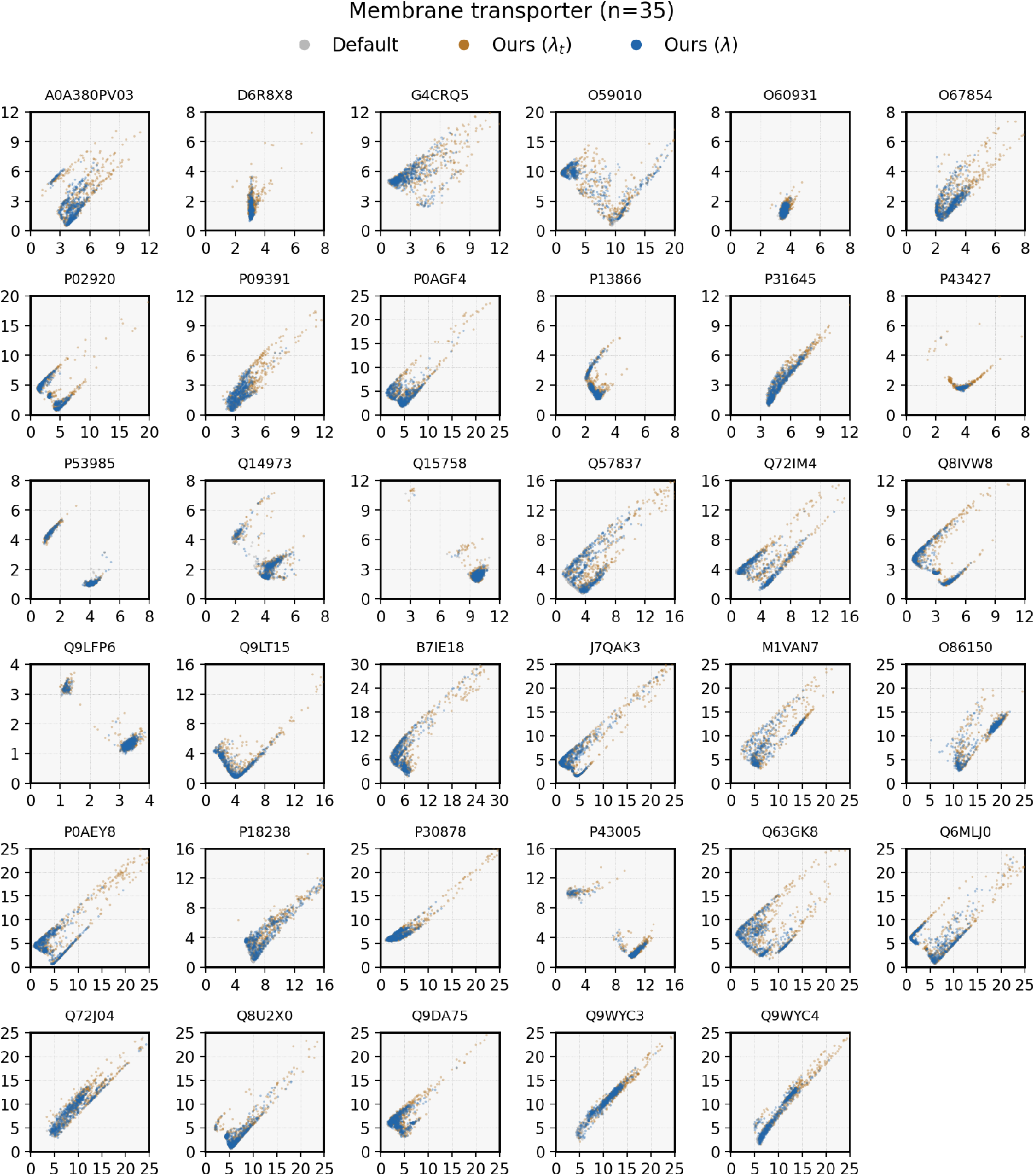
**Per-target 2D RMSD scatter for membrane transporters** (*n*=35).

**Figure 14.**
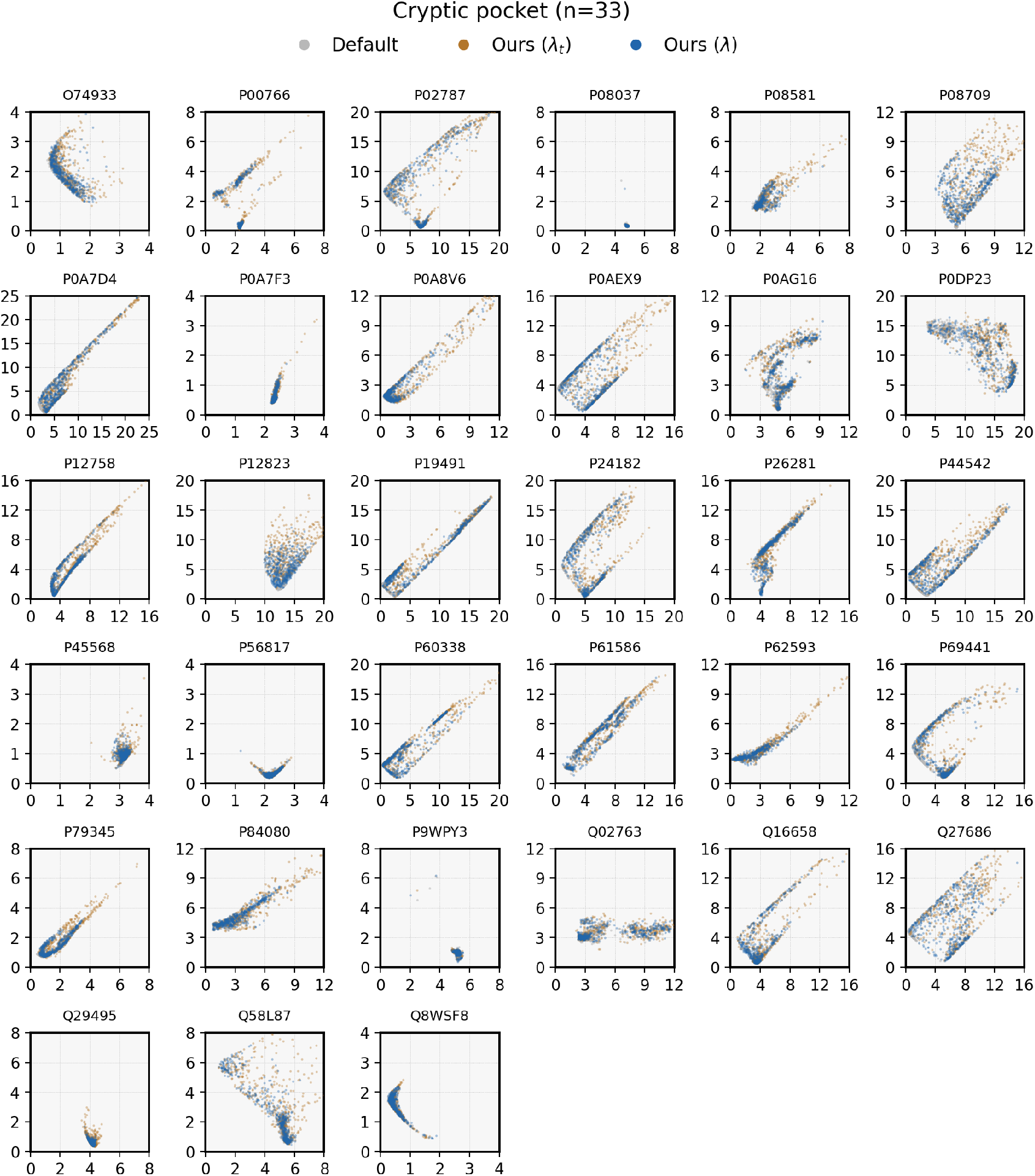
**Per-target 2D RMSD scatter for cryptic pockets** (*n*=33).

**Figure 15.**
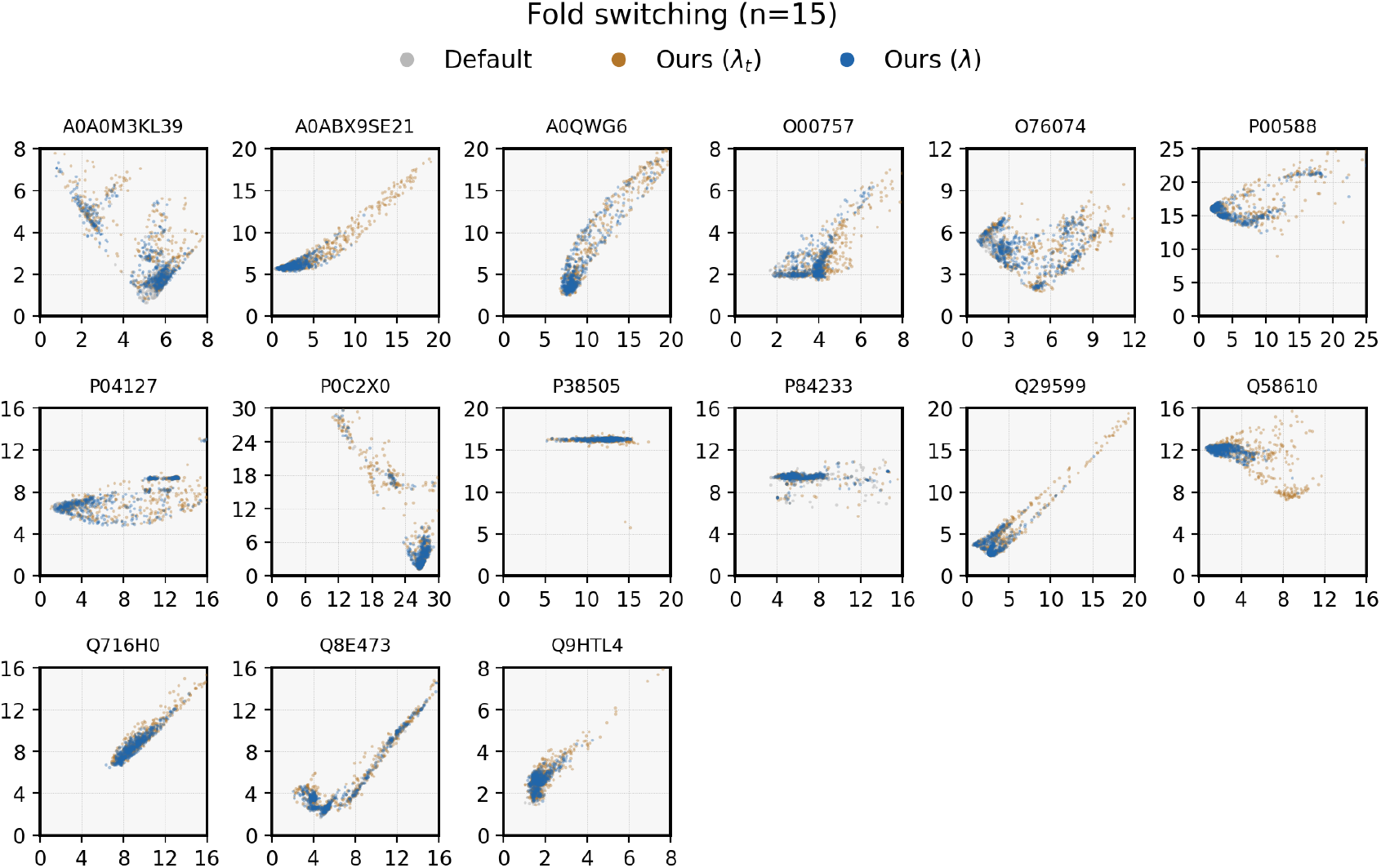
**Per-target 2D RMSD scatter for fold switching** (*n*=15).

**Figure 16.**
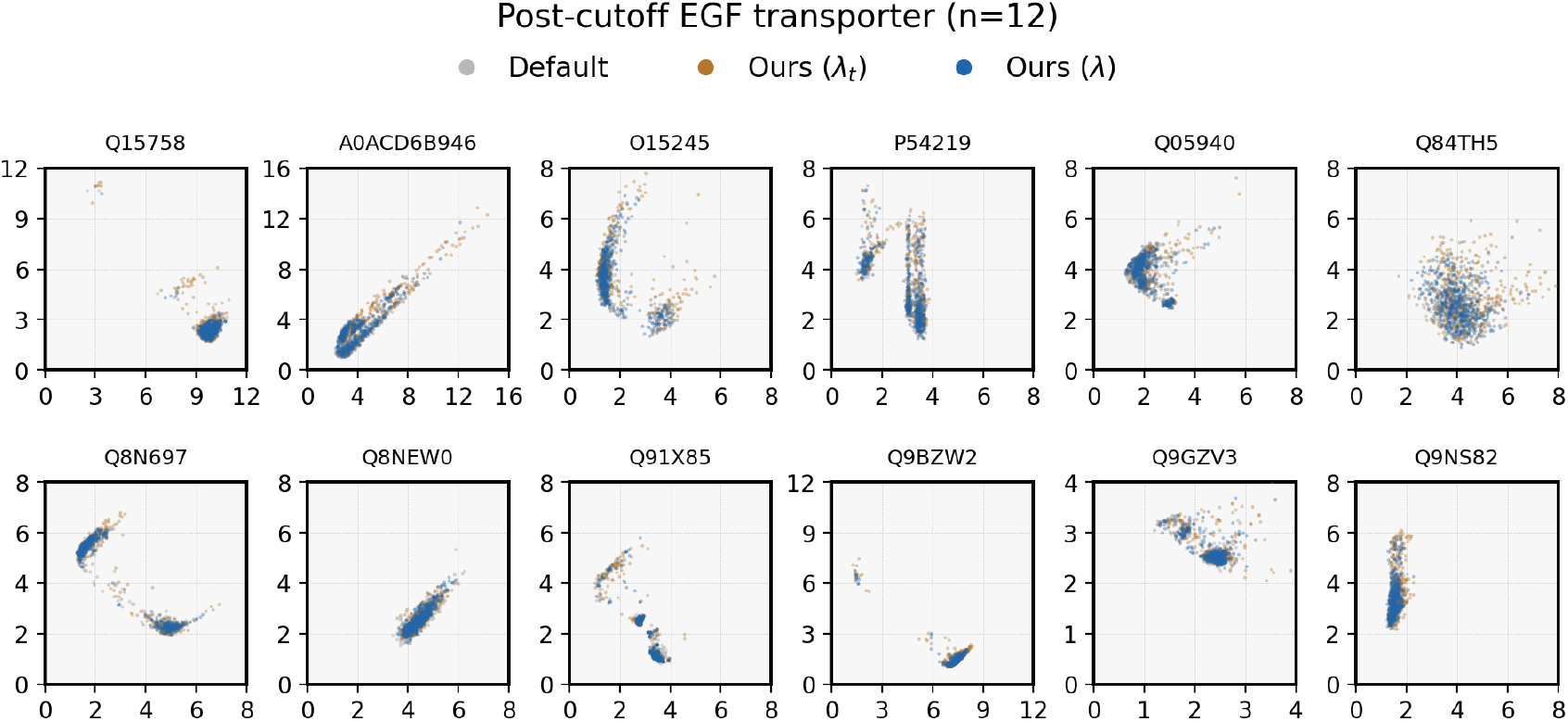
**Per-target 2D RMSD scatter for post-cutoff EGF transporters** (*n*=12). Targets match the Boltz-2 block of Table 4.

#### K.3 State classification and PCA

Each prediction is assigned to the state with the lowest global C*α* RMSD. The three candidate states are outward (8Y2G), occluded (8Y2D), and inward (minimum RMSD over 8Y2C, 8Y2E, and 8Y2F). A state assignment is labeled confident when the RMSD to the best-matching state is at least 0.3 Å lower than that to the second-best state. Otherwise, the prediction is assigned to a boundary class (Inward/Occluded, Occluded/Outward, or Inward/Outward). This assignment depends on the RMSD margin, independently of the model’s confidence score. 8Y2D and the nearest inward reference are 1.15 Å apart, within Boltz-2’s own prediction spread.

For the PCA visualization (Figure 4), we vectorize C*α*–C*α* pairwise distances on the 525 residues common to all five references and the predictions. We fit PCA to the prediction vectors retained by the confidence filter (Appendix D) and project the five reference vectors onto these axes. PC1 explains 62% of the variance and separates outward from inward, and PC2 explains 13% and separates occluded from the two open states.

#### K.4 Per-class results

Mean structural measurements by assigned state are reported in Table 11. The 43 ConforFlux predictions assigned to the inward-facing state have R85–D476 separation, F76–G425 cavity opening, and F76 displacement close to the experimental inward values, with a larger TM1a tilt. The boundary Inward/Occluded class (139 ConforFlux samples) is close to the inward references in C*α* RMSD but retains a narrower cytoplasmic cavity, consistent with incomplete gate opening.

#### K.5 Pre-cutoff SLC6 inward structures

Earlier inward SLC6 structures include the human serotonin transporter (PDB 6DZZ, released 2019-04-24) and bacterial LeuT (PDB 3TT3, released 2012-01-04), both before Boltz-2’s 2023-06-01 cutoff [Passaro et al., 2025]. The DAT evaluation tests coverage of newly released DAT structures within this established transporter cycle.

### L Per-target RMSD distributions

Each panel shows the C*α* Kabsch RMSD of every sample to the first reference against its RMSD to the second. Cryptic-pocket panels use the annotated pocket region. Default Boltz-2 and both settings of Eq. (2) are overlaid.

